# Small Glycols Discover Cryptic Pockets on Proteins for Fragment-based Approaches

**DOI:** 10.1101/605121

**Authors:** Harsh Bansia, Pranjal Mahanta, Neela H Yennawar, Suryanarayanarao Ramakumar

## Abstract

Cryptic pockets are visible in ligand-bound protein structures but are occluded in unbound structures. Utilizing these pockets in fragment-based drug-design provides an attractive option for proteins not tractable by classical binding sites. However, owing to their hidden nature, they are difficult to identify. Here, we show that small glycols find cryptic pockets on diverse set of proteins. Initial crystallography experiments serendipitously revealed the ability of ethylene glycol, a small glycol, to identify a cryptic pocket on W6A mutant of RBSX protein (RBSX-W6A). Explicit-solvent molecular dynamics (MD) simulations of RBSX-W6A with exposed-state of the cryptic pocket (ethylene glycol removed) revealed closure of the pocket reiterating that cryptic pockets in general prefer to stay in closed-state in absence of the ligands. Also, no change in the pocket was observed for simulations of RBSX-W6A with occluded-state of the cryptic pocket, suggesting that water molecules are not able to open the cryptic pocket. “Cryptic-pocket finding” potential of small glycols was then supported and generalized through additional crystallography experiments, explicit-cosolvent MD simulations, protein dataset construction and analysis. The cryptic pocket on RBSX-W6A was found again upon repeating the crystallography experiments with another small glycol, propylene glycol. Use of ethylene glycol as probe molecule in cosolvent MD simulations led to the enhanced sampling of the exposed-state of experimentally observed cryptic sites on test set of two proteins (Niemann-Pick C2, Interleukin-2). Further, analyses of protein structures with validated cryptic sites showed that ethylene glycol molecules binds to sites on proteins (G-actin, Myosin II, Bcl-xL, Glutamate receptor 2) which become apparent upon binding of biologically relevant ligands. Our study thus suggests potential application of the small glycols in experimental and computational fragment-based approaches to identify cryptic pockets in apparently undruggable and/or difficult targets, making these proteins amenable to drug-design strategies.

---

Use of structural information concerning binding site(s) on validated protein targets [1] is often the starting point of any drug-design process. However, many of the validated targets are not easily druggable owing to presence of undesirable traits in them such as featureless binding sites, lack of complementary hydrogen bonding partners [2, 3]. Adding to this notion of undruggability is the observation that many pharmaceutically important targets have cryptic binding sites [4, 5]. which are occluded in unbound proteins but become apparent in ligand-bound structures [6, 7] thereby making structural information available only on the unbound protein inadequate for drug-design purposes. Nonetheless, finding and targeting cryptic binding sites presents an attractive opportunity for many of the targets that are not tractable by traditional drug-design strategies, thereby expanding the druggable genome [8]. For years, efforts to develop inhibitors against K-RAS, an oncogene mutated in human cancers, were unsuccessful until a new cryptic site was found leading to successful targeting of K-RAS [4, 9], thus emphasizing the importance of finding cryptic sites in the course of drug discovery. Consequently, identifying and utilizing cryptic binding sites for therapy has gained momentum in the past few years [10]. However, identification of cryptic binding sites is a daunting task owing to their hidden nature and also because molecular mechanism(s) by which cryptic sites are formed are not properly understood [4].

Serendipity has accounted for revelation of cryptic sites in several protein structures [9, 11–13] enabling characterization of these sites which in turn has been helpful in developing methods for identification of cryptic sites. Current approaches to cryptic-site discovery include computational methods such as, CryptoSite [6], long-timescale molecular dynamics (MD) simulations combined with Markov state models [14, 15] or fragment probes [4, 5], mixed-solvent MD simulations [16], tools for mapping small-molecule binding hot spots [17] and experimental methods such as, extensive screening of small fragments [18], site-directed tethering [19, 20]. Although, CryptoSite, a cryptic site prediction tool based on machine learning and trained on a representative dataset of protein structures with validated cryptic sites, is a promising approach, it may be limited by the availability of experimentally determined cryptic sites which constitute the training set. The success of mixed-solvent MD simulations and hot spot mapping tools depends upon the nature of probe molecules used for cryptic site identification and are not always successful. Fragment screening experiments are expensive, time consuming and often have negative outcomes. Importantly, the information that, which of the probes used in fragment screening have potential to identify cryptic sites is lacking. The identity of such probe molecules having validated “cryptic-site finding” potential can significantly reduce time, efforts and expenditure in fragment screening experiments for identification of cryptic sites. If such probe molecules are also amenable to mixed-solvent MD simulations and hot spot mapping protocols, these computational methods incorporating the “cryptic-site finding” probe molecules can be used to identify cryptic sites on modeled structures of protein targets which are difficult to crystallize.

In the present work, we show that small glycols act as finders of cryptic pockets on proteins. Initial serendipitous observation made through crystallography experiments, conducted on a protein system (RBSX-W6A, Trp6 to Ala mutant of recombinant xylanase from *Bacillus sp. NG-27)* studied in our laboratory, showed the ability of ethylene glycol molecule to discover a cryptic pocket on RBSX-W6A. Explicit-solvent MD simulations of RBSX-W6A with exposed-state of the cryptic pocket revealed closure of the pocket in absence of glycol molecule whereas no change in the cryptic pocket was observed in MD simulations of RBSX-W6A with occluded-state of the cryptic pocket, showing that cryptic pockets are unstable without ligands and prefer to stay in closed state in their absence [4, 21]. Upon repeating the crystallography experiments with propylene glycol, another small glycol, the cryptic pocket of RBSX-W6A was rediscovered further supporting the finding that small glycols can discover cryptic pockets. The cryptic pocket of RBSX-W6A showed properties similar to those of cryptic sites characterized in other protein systems [6], and interacted with glycols through hydrogen bond and van der Waals contacts. The combined crystal structure and simulation results thus justify the role of glycols in identifying the cryptic pocket of RBSX-W6A. Further, by using ethylene glycol as a probe molecule in cosolvent MD simulations, we demonstrate its ability to induce opening of experimentally validated cryptic sites on test set of two proteins (Interleukin-2, Niemann-Pick C2). Finally, through protein dataset construction and analysis of protein structures with validated cryptic sites, we show that ethylene glycol molecules bind to cryptic sites in other proteins including targets of pharmaceutical interest for which the same cryptic sites become apparent upon binding of biologically relevant ligands. Our study thus validates and generalizes the “cryptic-pocket finding” potential of small glycols in proteins.

Use of cryptic binding sites for therapy depends crucially on their successful identification. Accordingly, small glycols can be used in both experimental as well as computational methods to identify cryptic pockets on proteins thereby facilitating fragment-based drug-design for apparently undruggable and/or difficult protein targets.

## Results and Discussion

### Ethylene glycol, a small glycol, identifies a cryptic pocket in RBSX-W6A crystal structure

During the course of studies aimed at characterizing the importance of aromatic residues in the stability of a recombinant xylanase from *Bacillus sp. NG-27* (RBSX) [22], we observed the presence of an ethylene glycol (1,2-ethanediol) (Ligand ID EDO) molecule occupying a surface pocket in the crystal structure of RBSX Trp6 to Ala mutant (RBSX-W6A) (PDB code 5EFD, Table S1) (Figure 1A), EDO was used as a cryoprotectant in the diffraction experiment (Methods, Supporting Information) and was clearly detected in *2mFo-DFc* electron density map contoured at l.0*σ*-level (Figure 1B). It must be noted that this surface pocket was not observed in the native RBSX structure (PDB code 4QCE) [23]. Analysis of atoms lining the surface pocket in 5EFD (chain A) and their atom-wise solvent accessibilities (AwSA (Å^2^)) revealed that this surface pocket is composed of both polar and apolar atoms (Table S2), Further, the EDO molecule interacts with the pocket atoms by forming a hydrogen bond (3.0 Å) with a polar, solvent accessible (AwSA: 1,059 Å^2^), backbone amide nitrogen atom of Ala6 along with a number of van der Waals (Vdw) contacts (18, d ≤ 4Å) (Figure S1A), Studies have shown that interaction of surface water also called as hydration water, with the protein surface affects protein structure, stability and function [24–27], The above observations led us to wonder if in the absence of the EDO molecule, a water molecule would interact with and occupy the amphiphilic surface pocket found in 5EFD, prompting us to redetermine the crystal structure of RBSX-W6A without the use of cryo-protectant ethylene glycol in the diffraction experiment (PDB code 5XC0, Table S1), Much to our surprise, the particular surface pocket observed in 5EFD was not seen in 5XC0 and the backbone amide nitrogen atom of Ala6 was also found to be solvent inaccessible (AwSA:0.000 Å^2^).

**Figure 1:**
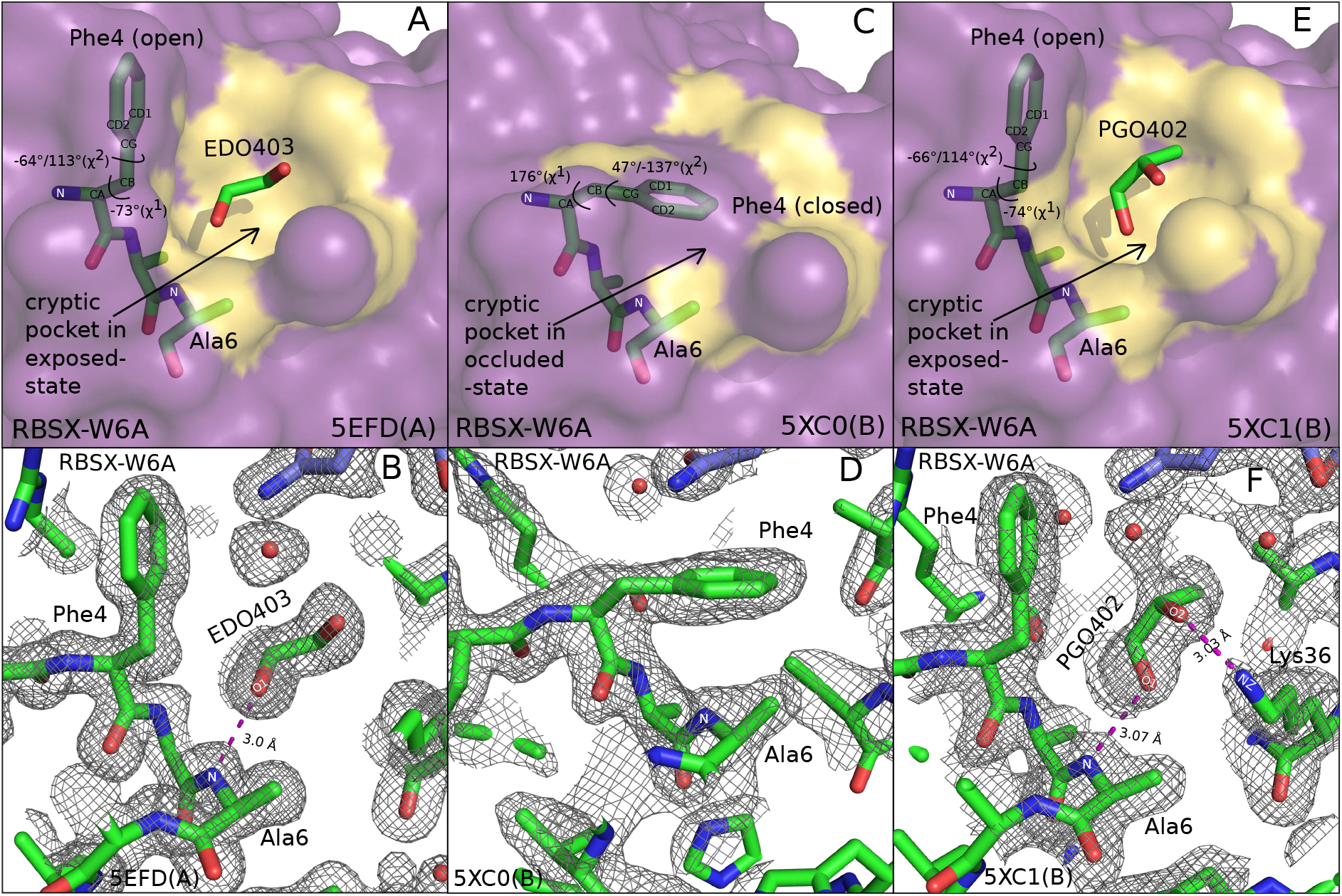
Small glycols, ethylene glycol and propylene glycol, identify a cryptic pocket on RBSX-W6A. Amino acid residues (green) and bound ligands (green) are shown in a stick model. Nitrogen and oxygen atoms are colored blue and red respectively while carbon atoms are colored according to the color mentioned for residues or ligands. Hydrogen bonds are represented as dashed lines (magenta) labeled with the donor-acceptor distance in A. Surface of RBSX-W6A (violetpurple) is drawn and is superposed with its residue Phe4 to illustrate the open/closed conformation of Phe4 and the associated exposed-state/occluded-state of the cryptic surface pocket (lightorange) caused by the presence/absence of small glycols. The open and closed conformations of Phe4 are defined here in terms of its side chain *χ*^1^ and *χ*^2^ (measured along N,CA,CB,CG and CA,CB,CG,CD1/CD2 atoms respectively) dihedral angles. (A) Exposed-state of the cryptic surface pocket containing a molecule of EDO with Phe4 in open state is shown for RBSX-W6A (5EFD, chain A) when it was crystallized in the presence of ethylene glycol. (C) Occluded-state of the cryptic surface pocket with Phe4 in closed state is shown for RBSX-W6A (5XC0, chain B) when it was crystallized in the absence of ethylene glycol. (E) Exposed-state of the cryptic surface pocket containing a molecule of PGO with Phe4 in open state is shown for RBSX-W6A (5XC1, chain B) when it was crystallized in the presence of propylene glycol. (B),(D) and (F) Electron density maps (grey) corresponding to (A),(C) and (E) are shown. The *2mFo-DFc* electron density maps contoured at 1*σ*-level show well defined electron density for EDO and PGO in (B) and (F) respectively.

Further analysis revealed that in the absence of the EDO molecule, the aromatic side-chain of the neighboring Phe4 residue moves and covers the surface pocket (Figure 1C and D), occluding it and thereby making backbone amide nitrogen atom of Ala6 solvent inaccessible. Based on the occluded-state and exposed-state of this surface pocket in 5XC0 and 5EFD respectively, we term it as a cryptic surface pocket which, otherwise masked by the side-chain of Phe4 residue, becomes apparent only when an EDO molecule binds to it. Residue Phe4 is observed to have two distinct conformations in 5XC0 and 5EFD, characterized by its side-chain *χ*^1^ (measured along N,CA,CB and CG atoms) and *χ*^2^ (measured along CA,CB,CG and CD1/CD2 atoms) dihedral angles. We term the conformation of Phe4 in 5XC0 as “closed state” (*χ*^1^= 176°, *χ*^2^= 47°/−137°) (Figure 1C), where it occludes the cryptic surface pocket in absence of the EDO molecule, and in 5EFD as “open state” (*χ*^1^= −73°, *χ*^2^= −64°/113°) (Figure 1A), where it gets displaced exposing the cryptic surface pocket upon binding of the EDO molecule. Further, a measurement of the solution affinity constant between ethylene glycol molecules and RBSX/RBSX-W6A was performed using isothermal titration calorimetry (ITC) experiments (Methods, Supporting Information), No measurable binding was seen between ethylene glycol and the RBSX while weak binding was detected for the RBSX-W6A (Figure S2, Table S3) suggesting binding of ethylene glycol molecules to RBSX-W6A in solution as well under physiological conditions. It must be noted that small molecular probes binding with weak affinity such as ethylene glycol may be inaccessible to some of the current biophysical methods of detection [28], Also, compared to other methods, X-ray crystallography can detect binding of MiniFrags, ultra-low molecular weight compounds binding with very low affinities [29], an observation reiterated here through crystallography for binding of ethylene glycol to RBSX-W6A.

### Explicit-solvent MD simulations of RBSX-W6A reveals closure of the cryptic pocket in absence of ethylene glycol molecule

We further computationally explored the possibility whether water molecules, in a way similar to an EDO molecule, can induce the transition of Phe4 from closed to open state and the associated opening of the cryptic surface pocket. For this, we conducted an all-atom explicit-solvent molecular dynamics (MD) simulations starting with RBSX-W6A structure (5XC0, chain B) in which Phe4 was observed in the closed state occluding the cryptic surface pocket (Methods, Supporting Information). The side-chain *χ*^1^ and *χ*^2^ dihedral angles for the Phe4 residue and radial distribution function of water around backbone amide nitrogen atom of Ala6, as indicators of transition of Phe4 from closed to open state and the corresponding transition of cryptic surface pocket from occluded-state to exposed-state, were monitored from the simulation trajectories. The peaks in radial distribution function of water (represented as g(r), where r stands for the distance from the selected atoms), represent the hydration shells around the selected atoms. The population distribution of Phe4 measured every 30 ns in terms of its side-chains *χ*^1^ and *χ*^2^ dihedral angles from the simulation trajectories, revealed that the Phe4 remained in the closed state and therefore the cryptic surface pocket remained in the occluded-state throughout the length (120 ns) of the MD simulations with the most populated states of Phe4 being defined by *χ*^1^ = −170° to 160° (centered at 180°) and *χ*^2^ = −140° to −100° (centered at −120°) (Figure 2A-D). Interestingly, upon conducting all-atom explicit-solvent MD simulations starting with RBSX-W6A structure (5EFD, chain A, EDO removed) where Phe4 is present in open state exposing the cryptic surface pocket (Methods, Supporting Information), similar analysis on the simulation trajectories showed that Phe4 stayed in the open state and therefore the cryptic surface pocket remained in exposed-state for approximately first 30 ns, *χ*^1^ = −70° to −50° (centered at −60°) and *χ*^2^ = −80° to −50° (centered at −70°), (Figure 2E, 1-30 ns). Thereafter Phe4 started transitioning to closed state with the cryptic surface pocket transitioning to occluded state (Figure 2F, 31-60 ns). After that Phe4 stayed closed and therefore the cryptic surface pocket stayed occluded for the remaining simulation time, *χ*^1^ = −170° to 160° (centered at 180°) and *χ*^2^ = −130° to −90° (centered at −110°) (Figure 2G and H, 61-120 ns).

**Figure 2:**
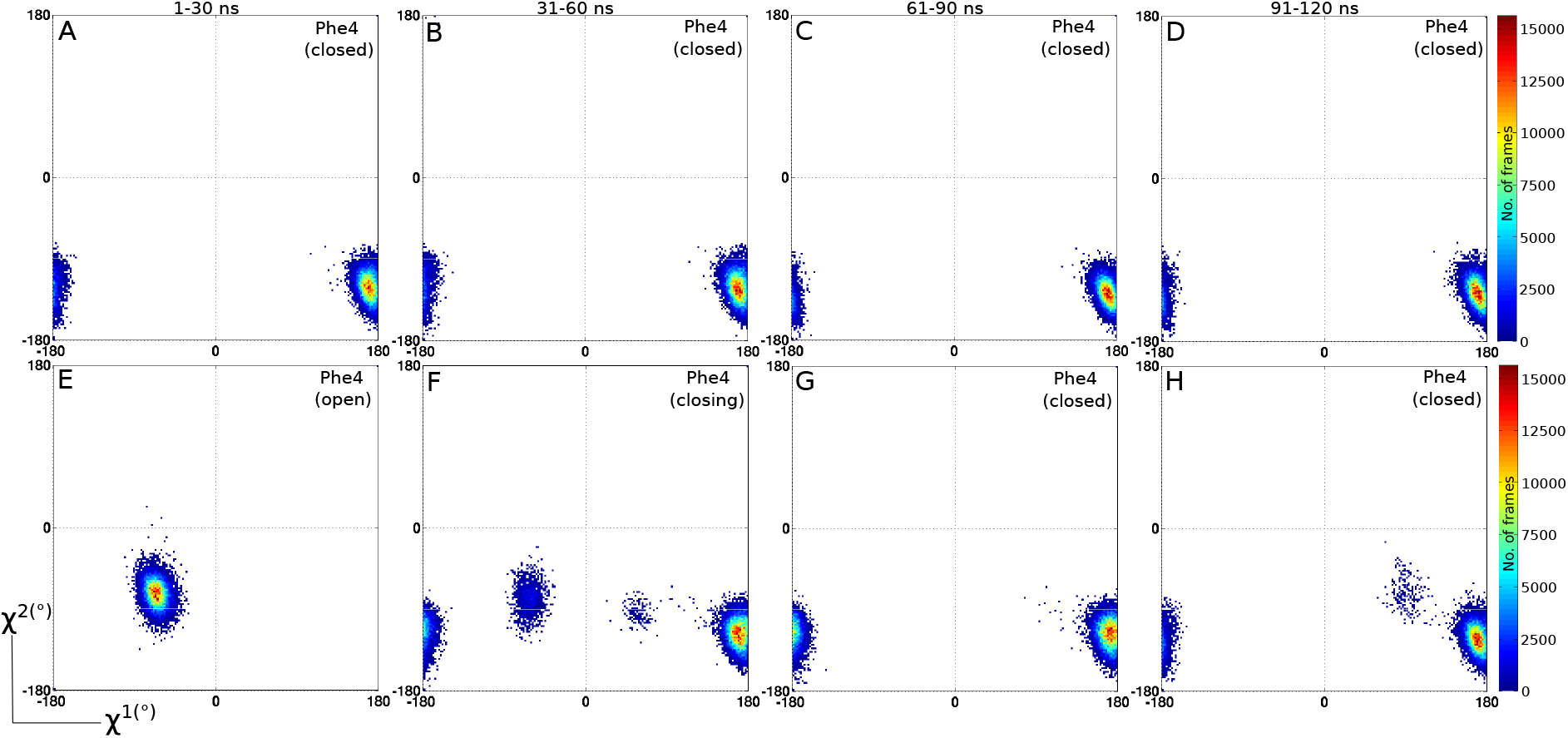
2D-histogram plots showing population distribution of residue Phe4 between closed and open states during 120 ns long explicit-solvent MD simulations of RBSX-W6A. The distribution measured every 30 ns in terms of *χ*^1^ and *χ*^2^ side-chain dihedral angles of residue Phe4 shows that for the entire simulation time (A)-(D) Phe4 remains in closed state and therefore the cryptic surface pocket remains in occluded-state when RBSX-W6A structure (5XC0, chain B) with Phe4 in closed state occluding the cryptic surface pocket was used to initialize the explicit-solvent MD simulations. For explicit-solvent MD simulations, started with RBSX-W6A structure (5EFD, chain A, EDO removed) in which Phe4 is in open state exposing the cryptic surface pocket, the same analysis shows that (E) for approximately first 30 ns Phe4 remains in open state and therefore the cryptic surface pocket remains in exposed-state, then (F) Phe4 transitions to closed state with the cryptic surface pocket transitioning to occluded-state and after that (G), (H) Phe4 stays closed for rest of the simulation and the cryptic surface pocket remains occluded.

Further, the radial distribution function of water calculated from the simulation trajectories of 5EFD (chain A) reveals that the hydration shell which occurs at 2.95 Å from backbone amide nitrogen atom of Ala6 when Phe4 is in open state exposing the cryptic surface pocket (1-30 ns) (Figure 3B), disappears when Phe4 transitions from open to closed state occluding the cryptic surface pocket. Thereafter (31-120 ns) (Figure 3B) the radial distribution function of water in MD simulations of 5EFD (chain A) resembles to the radial distribution function of water calculated from the simulation trajectories of 5XC0 (chain B) (Figure 3A), reiterating observation made for crystal structure 5XC0 that backbone amide nitrogen atom of Ala6 is solvent inaccessible when Phe4 is in the closed state occluding the cryptic surface pocket.

**Figure 3:**
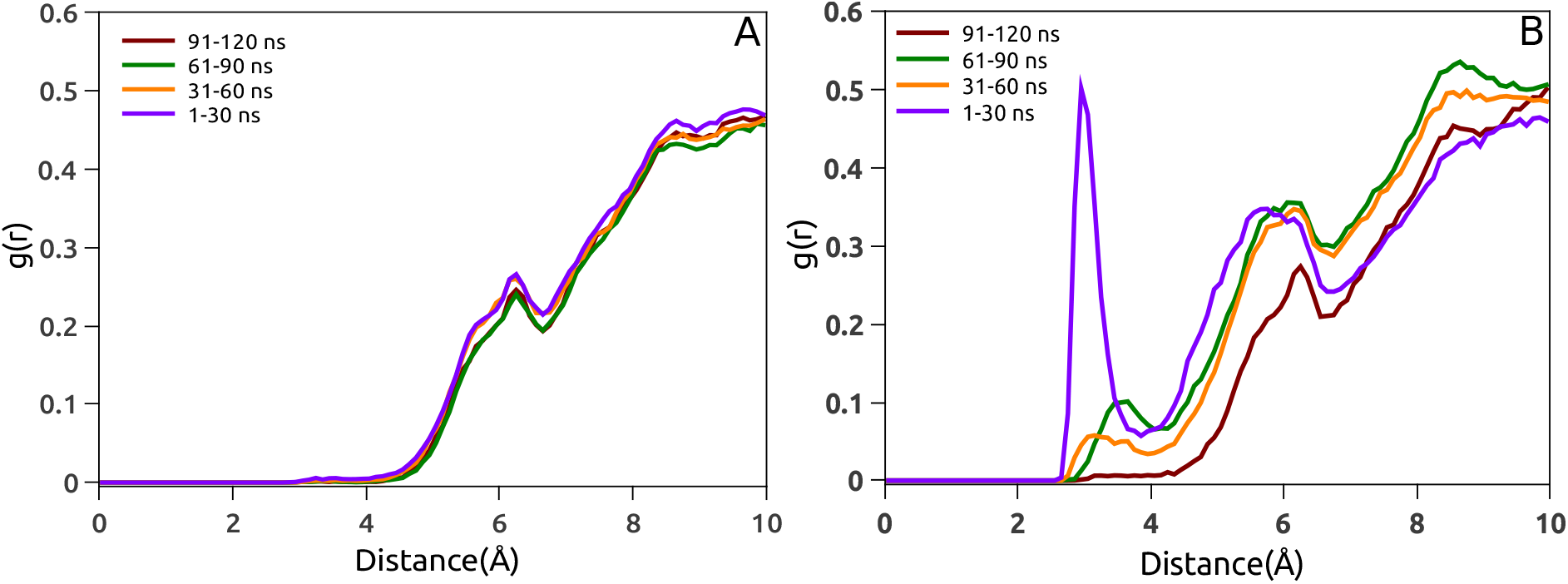
Radial distribution function of water around the backbone amide nitrogen atom of Ala6 calculated from the MD simulation trajectories of RBSX-W6A. The peaks in radial distribution function of water (represented as g(r), where r stands for the distance from the selected atoms), represent the hydration shells around the selected atoms and for RBSX-W6A shows that (B) the hydration shell, which occurs at 2.95 Å from the backbone amide nitrogen atom of Ala6 when Phe4 is in open state exposing the cryptic surface pocket (1-30 ns), is not observed in (A) 1-120 ns and(B) 31-120 ns, indicating that backbone amide nitrogen atom of Ala6 is solvent inaccessible when Phe4 is in closed state occluding the cryptic surface pocket.

Thus the simulation results support our experimental findings that in the hydrated environment without EDO molecules, the closed state of Phe4 occluding the cryptic surface pocket is preferred and the presence of EDO molecule stabilizes the open state of Phe4 exposing the cryptic surface pocket. It has been shown also, through use of various force-field in MD simulations of different protein systems with known cryptic sites, that cryptic pockets are unstable without ligands and prefer to stay in closed state in their absence [4, 21].

### Propylene glycol, another small glycol identifies the cryptic pocket in RBSX-W6A crystal structure

Based on the results obtained so far, we hypothesized that small glycols can discover the cryptic surface pocket of RBSX-W6A by displacing the aromatic side-chain of Phe4 residue. To test the hypothesis, we redetermined the crystal structure of RBSX-W6A in the presence of another small glycol, propylene glycol (Propane-1,2-diol, racemic mixture) (PDB code 5XC1, Table S1) and reexamined the status of the cryptic surface pocket. As anticipated, we re-observed the cryptic surface pocket in exposed-state, this time containing a molecule of S-1,2-propanediol (Ligand ID PGO) with the Phe4 in the open state (*χ*^1^= −74°, *χ*^2^= −66°/114°) (Figure 1E), thus vindicating our hypothesis. The cryptic surface pocket in 5XC1 shares similar attributes with the cryptic surface pocket in 5EFD (Table S2), Similar to the scenario observed for the EDO molecule in 5EFD, in 5XC1 also, the identified cryptic pocket was found to be amphiphilic in nature (Table S2) and the PGO molecule was observed to participate in hydrogen bonding interaction (3.07 Å) with the solvent accessible polar backbone amide nitrogen atom of Ala6 (AwSA:l.393 Å^2^) which has unsatisfied hydrogen bonding potential (Figure S1B). PGO in the pocket also hydrogen bonds with Lys36 (Figure S1B) and forms a number of van der Waals (Vdw) contacts (21, d ≤ 4 Å) with the pocket atoms. A recent comprehensive study, characterizing cryptic sites in different protein systems, has also reported that cryptic sites tend to be less hydrophobic and more flexible [6]. The finding that these small glycols (EDO and PGO) identify and interact with the cryptic surface pocket in a similar manner suggests a certain level of selectivity to their mode of interaction with this pocket. Moreover in 5XC1, of the several surface pockets, we were able to model PGO (*2mFo-DFc* electron density map (Figure 1F)) only in this particular surface pocket emphasizing the non-random nature of the interaction. It is interesting to note that the rotational freedom about the carbon-carbon single bond enables small glycols, EDO and PGO, to adopt different conformations out of which these glycols display amphiphilic character in gauche^+^ and gauche^−^ conformations. Accordingly, the inherent flexibility, amphiphilicity of small glycols would enable an interplay between the dynamics of the flexible, less hydrophobic cryptic binding site and the dynamics of the glycol molecule interacting with the site. Although not observed in 5XC1, we cannot rule out the possibility of R-1,2-propanediol (Ligand ID PGR) acting in a way similar to that of PGO and EDO. We summarize that in RBSX-W6A, the substitution of bulky Trp6 to Ala creates a surface pocket that remains occluded by the aromatic side-chain of Phe4 residue as observed in 5XC0. When EDO/PGO molecules are allowed to interact with RBSX-W6A, they not only displace the aromatic side-chain of Phe4 exposing the cryptic surface pocket but also interact with the pocket in a similar manner as seen in 5EFD and 5XC1. Thus, the combined crystal structure and simulation results justifies the role of small glycols in identifying the cryptic pocket of RBSX-W6A. Importantly, cryptic cavity/cavity generated on the protein surface at the mutation site has been successfully used to target the oncogenic mutants, K-Ras(G12C) [9] and p53(Y220C) [30], assuming that binding of small molecules, while inhibiting the oncogenic mutant, would not affect the functioning of the wild-type protein lacking the mutational cavity. This suggests that cryptic cavities/cavities arising due to mutations are equally important for therapy in addition to such sites inherently present in wild-type proteins. Based on our work it may be surmised that determination of protein crystal structures in the presence and absence of small glycols (ethylene glycol and propylene glycol) and a systematic comparison of the protein structures could help in the identification of cryptic sites in the protein under study.

### Enhanced sampling of the exposed-state of experimentally validated cryptic sites of other proteins by ethylene glycol molecules in explicit-cosolvent MD simulations

Our so far discussed results concluded from the studies conducted on RBSX-W6A, used as a model protein, are thus suggestive of a role of the small glycols in finding cryptic pockets on proteins. Motivated by the application of cosolvent MD simulations in identifying cryptic sites [16, 31], we wondered whether the use of ethylene glycol as cosolvent in MD simulations could unveil cryptic sites on other proteins. A variety of small molecules containing aromatic and aliphatic moieties substituted with different chemical functional groups, have been used as probes in cosolvent MD simulations [32]. However, it must be important to note that the use of ethylene glycol as cosolvent in MD simulations to identify binding hotspots and/or cryptic sites has not been reported earlier to the best of our knowledge. Over the past few years, cosolvent-based MD simulations approach has been increasingly employed to assess druggability of targets of pharmaceutical interest by identifying binding sites and hotspots on proteins [33] and has also shown to enhance the sampling of exposed states of cryptic sites on a range of protein targets [16, 31]. We chose two proteins, Niemann-Pick C2 (NPC-2) and Interleukin-2 (IL-2), for ethylene glycol based cosolvent MD simulations. The choice was motivated by the observation that these proteins are pharmaceutically important, harbor experimentally validated cryptic sites and have been previously subjected to cosolvent MD simulations using different probes (other than ethylene glycol) in order to identify druggable cryptic binding sites [16, 31].

Most importantly, to the best of our knowledge small glycols studied in this report have not been used in the crystallization experiments of NPC-2 and IL-2. For cosolvent MD simulations of NPC-2 and IL-2, we used a 5 % v/v ethylene glycol-water solution. Use of high cosolvent concentrations (15 % or above) have been shown to induce protein unfolding and therefore are discouraged [31]. Also, a 5% concentration is close to concentrations used in experiments [34]. Details of system preparation and subsequent simulations of NPC-2 and IL-2 for ethylene glycol based cosolvent MD simulations is given in Methods, Supporting information. Here, we present the results of cosolvent simulations from the point of view of potential of ethylene glycol molecule(s) to unmask the experimentally validated cryptic sites of NPC-2 and IL-2.

#### Niemann-Pick C2

(NPC-2) is a small protein involved in transport of cholesterol and other sterols from lysosomes to different cellular locations [35]. The deficiency of NPC-2 causes fatal Niemann-Pick type C2 disease characterized by accumulation of cholesterol in lysosomes [36]. In the holo-form of the NPC-2 structure (PDB ID 2HKA, Chain C) (Figure 4B), binding of cholesterol-3-O-sulfate (Ligand ID C3S) disrupts the *π*-stacking interaction and displaces side-chains of residues, Phe66 on the *βD* strand and Tyr100 on the *βE-βF* loop, thereby revealing a deep cavity not observed in the apo-form of the NPC-2 structure (PDB ID 1NEP, Chain A) (Figure 4A) [37]. Thus, the cryptic site on NPC-2 structure is an orthosteric site and represents a challenging test case for our ethylene glycol based cosolvent simulations due to the buried hydrophobic nature of the cryptic pocket [5]. Apo-form of the NPC-2 structure was subjected to short (50 ns) explicit-solvent and cosolvent MD simulations with ethylene glycol as probe molecules (Methods, Supporting information) to see whether ethylene glycol molecules could enhance sampling of the open-state of the NPC-2 cryptic pocket. Comparison of apo and holo structures of NPC-2 shows that binding of cholesterol-3-O-sulfate to NPC-2 and opening of the cryptic pocket is accompanied by a significant change in *χ*^1^ (measured along N,CA,CB,CG atoms) dihedral angle of Phe66 from −90.8° in apo NPC-2 structure (Figure 4A) to −175.4° in holo NPC-2 structure (Figure 4B) and displacement of residues Phe66, Tyr100 as measured here by increase in the distance between CA atoms of residues Phe66 and Tyr100 from 9.1Å in apo NPC-2 structure (Figure 4A) to 12.7 Å in holo NPC-2 structure (Figure 4B). Comparison of apo and holo structures of NPC-2 also shows that there is no significant change in *χ*^1^ dihedral angle of Tyr100(−56.3° in apo NPC-2 structure, Figure 4A and −65.1° in holo NPC-2 structure, Figure 4B). Therefore, transition in Phe66 *χ*^1^ side-chain dihedral angle and distance between CA atoms of residues Phe66 and Tyr100, as indicators of the dynamics of the cryptic orthosteric site, were monitored from the explicit-solvent and cosolvent simulation trajectories of NPC-2. Time series plots shows that, in comparison to the explicit-solvent simulation of NPC-2 (Figure 4C and 5A), presence of ethylene glycol molecules in cosolvent simulation considerably shifts/increases the distribution of Phe66 *χ*^1^ side-chain dihedral angle and distance between CA atoms of residues Phe66 and Tyr100 more towards the values defining the exposed-state of the NPC-2 cryptic pocket (Figure 4D and 5B). Representative structural snapshots sampled from the cosolvent simulation trajectory at 23^rd^ (Figure 4E) and 37^th^ ns (Figure 4F) depicts that ingression of cosolvent, ethylene glycol molecules, at the cryptic site causes residues, Phe66 and Y100, to assume conformations (23^rd^ ns: Phe66 *χ*^1^ −154.5°, Tyr100 *χ*^1^ −44,4°, Phe66CA-Tyr100CA distance 11.2 Å, Figure 4E; 37^th^ ns: Phe66 *χ*^1^ −151.0°, Tyr100 *χ*^1^ −50.0°, Phe66CA-Tyr100CA distance 11.0 Å, Figure 4F) similar to the site-permissible conformations of these residues (Phe66 *χ*^1^ −175.4Å, Tyr100 *χ*^1^ −65.1Å Phe66CA-Tyr100CA distance 12.7 Å, Figure 4B) observed in the holo-form of the NPC-2 structure, thereby identifying the cryptic orthosteric site (Figure 4E and F). Previous cosolvent MD simulation study of NPC-2 in presence of 5 % v/v isopropyl alcohol only explored regions intermediate to the closed and open forms whereas 5 % resorcinol based cosolvent simulation initially sampled exposed-state of the cryptic pocket but later on drove the sampling away from the holo structure [16]. Our 5% ethylene glycol based cosolvent simulation of NPC-2 echoes results similar to 5% ethylene glycol based cosolvent simulation of NPC-2 echoes results similar to 5% resorcinol wherein the distribution, though considerably shifted towards the open state in comparison to water only simulations of NPC-2 (Figure 4C and 5A), toggles back and forth between open and closed states of the cryptic pocket (Figure 4D and 5B) and might point to the limitation of smaller probe molecules in sampling cryptic sites revealed by binding of huge ligands. However, in spite of the size and chemical nature of ethylene glycol similar to isopropyl alcohol, it fared better than isopropyl alcohol and similar to bulkier aromatic resorcinol in cosolvent studies to identify cryptic pocket of NPC-2. The comparison made is only qualitative and not quantitative owing to the variations in cosolvent simulation methodology. In conclusion it may be seen that ethylene glycol molecules exposes the cryptic orthosteric site on NPC-2, also sampled by the aromatic probe, resorcinol [16].

**Figure 4:**
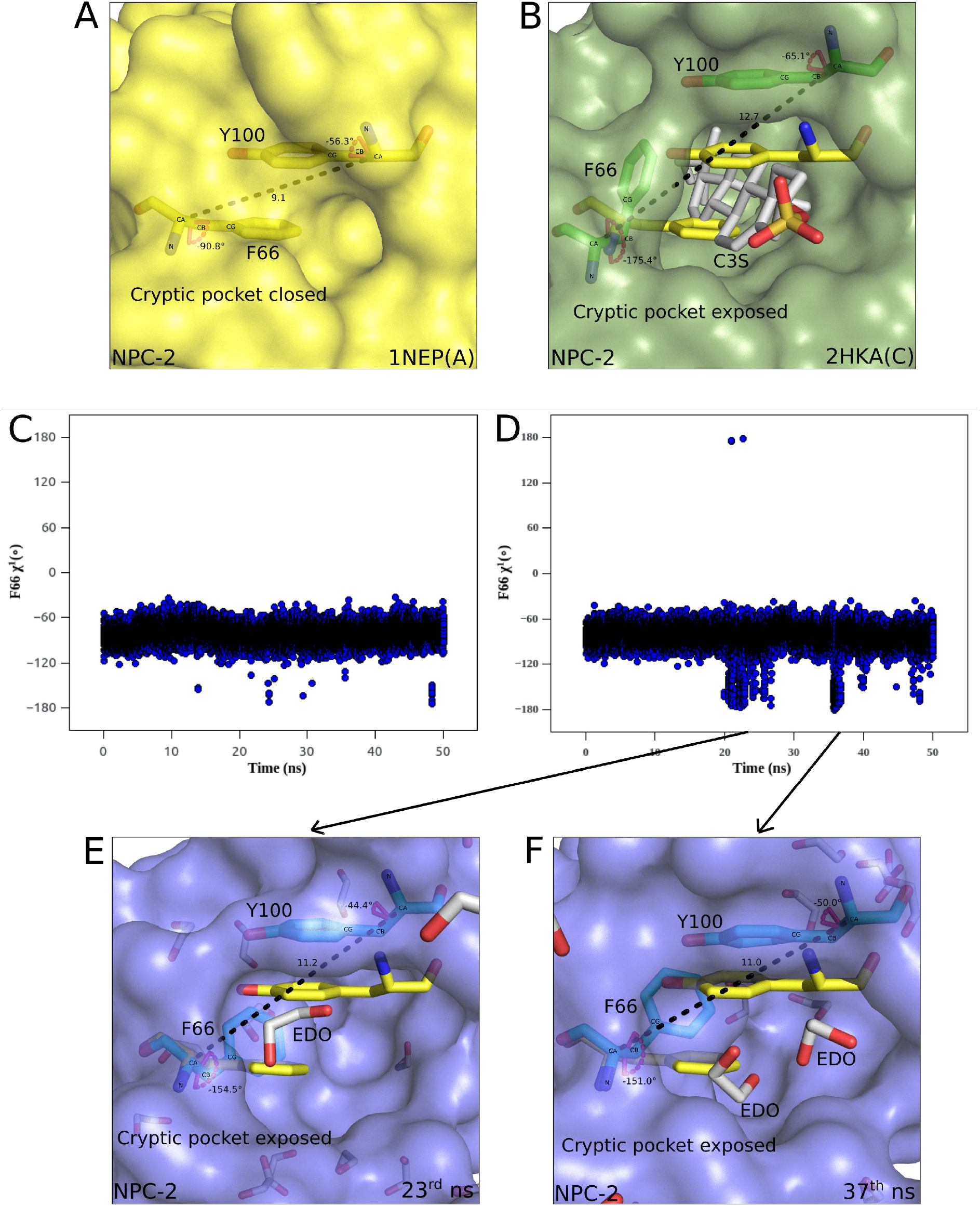
Enhanced sampling of the exposed-state of the cryptic pocket in ethylene glycol based cosolvent simulations of NPC-2 / Ethylene glycol based cosolvent simulations enhances sampling of the exposed-state of the cryptic pocket in NPC-2. Amino acid residues and bound ligands are shown in a stick model. Nitrogen, oxygen and sulphur atoms are colored blue, red and orange respectively while carbon atoms are colored according to the color mentioned for residues or ligands. Residue side-chain *χ*^1^ (measured along N,CA,CB,CG atoms) dihedral angles are shown as labeled magenta arcs. Residue displacements are represented as black dashed lines labeled with distance in Å. Surface of NPC-2 is drawn and is superposed with its residues Phe66, Tyr100 to illustrate the rearrangements of these residues and the associated opening of the cryptic pocket upon ligand binding. (A) Closed-state of cryptic pocket and associated arrangement of residues Phe66, Tyr100 (yellow sticks) is shown in unbound NPC-2 structure (yellow, 1NEP(A)). (B) Rearrangements of residues Phe66, Tyr100 (protruding, yellow sticks; displaced, green sticks) and opening up of the cryptic pocket upon binding of cholesterol-3-O-sulfate (Ligand ID C3S, grey) to NPC-2 structure (smudge, 2HKA(C)) is shown. Time series plots showing distribution of the *χ*^1^ side-chain dihedral angle of Phe66, as one of the parameters referring/indicating to the distribution of the closed/open states of the cryptic pocket, in (C) explicit-solvent and (D) 5 % ethylene glycol based explicit-cosolvent MD simulations of apo NPC-2 structure (1NEP(A)). Snapshots sampled/selected/extracted from the cosolvent simulation trajectory at (E) 23^rd^ ns and (F) 37^th^ ns depicts/shows rearrangements of residues Phe66, Tyr100 (protruding, yellow sticks; displaced, blue sticks) and associated opening up of the cryptic pocket upon binding of ethylene glycol molecule(s) (Ligand ID EDO, grey) to NPC-2 structure (slate).

**Figure 5:**
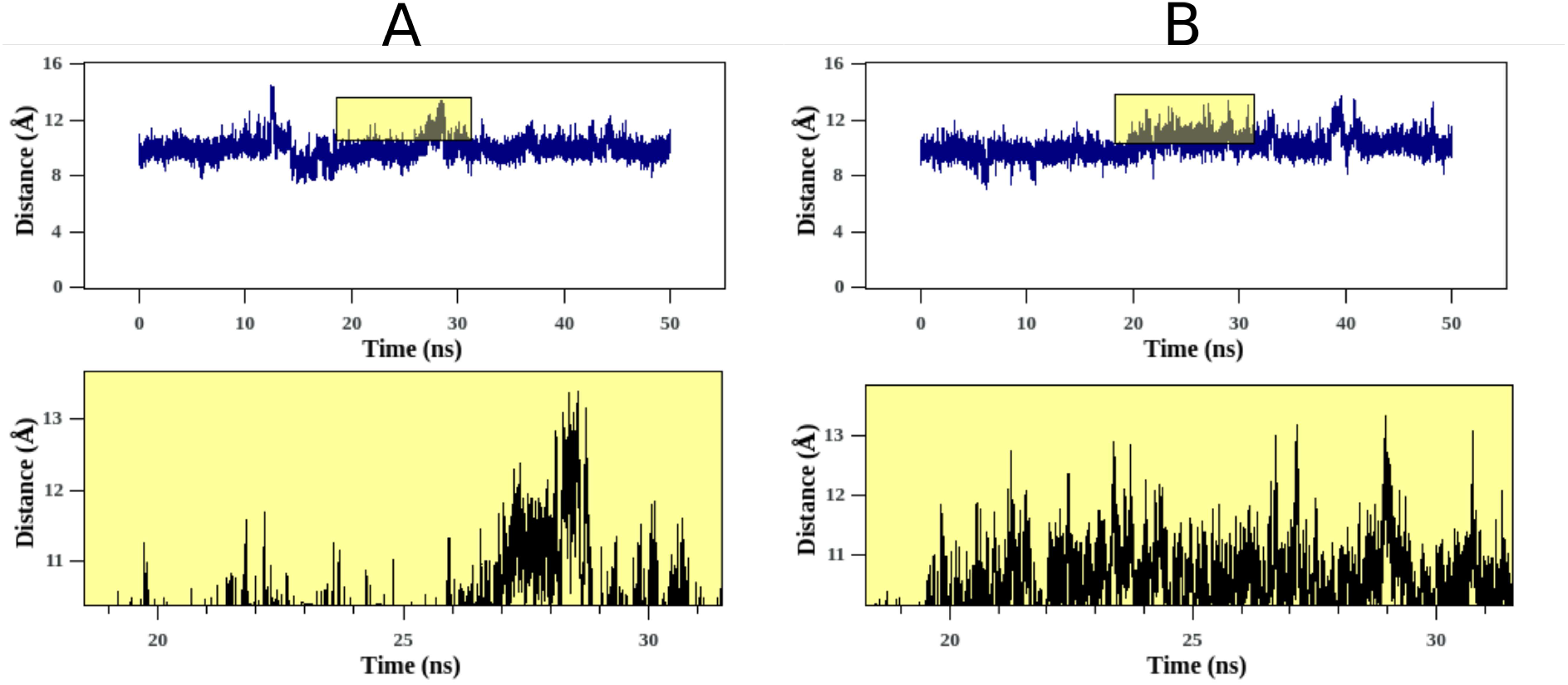
Time series plots showing distribution of the distance between CA atoms of residues Phe66 and Tyr100, as one of the parameters referring/indicating to the distribution of the closed/open states of the cryptic pocket, in (A) explicit-solvent and (B) 5 % ethylene glycol based explicit-cosolvent MD simulations of apo NPC-2 structure (1NEP(A)). Plots are zoomed at specific time-intervals to highlight the transition.

#### Interleukin-2

(IL-2) is a cytokine that acts by binding to various IL-2 receptors (IL-2Rs) [38]. It is of considerable pharmaceutical interest as it plays a significant role in the activation of T-cells and rejection of tissue grafts [39]. The cryptic site on IL-2 is located at the IL-2R*α* binding site and is occluded by the protrusion of side-chain of residue Phe42 in the apo-structure (PDB ID 1M47) (Figure 6A). Binding of a small molecule (Ligand IDs FRG) displaces the Phe42 side-chain thereby revealing the cryptic site in the holo-structure of IL-2 (PDB IDs 1M48) (Figure 6B) [40].

**Figure 6:**
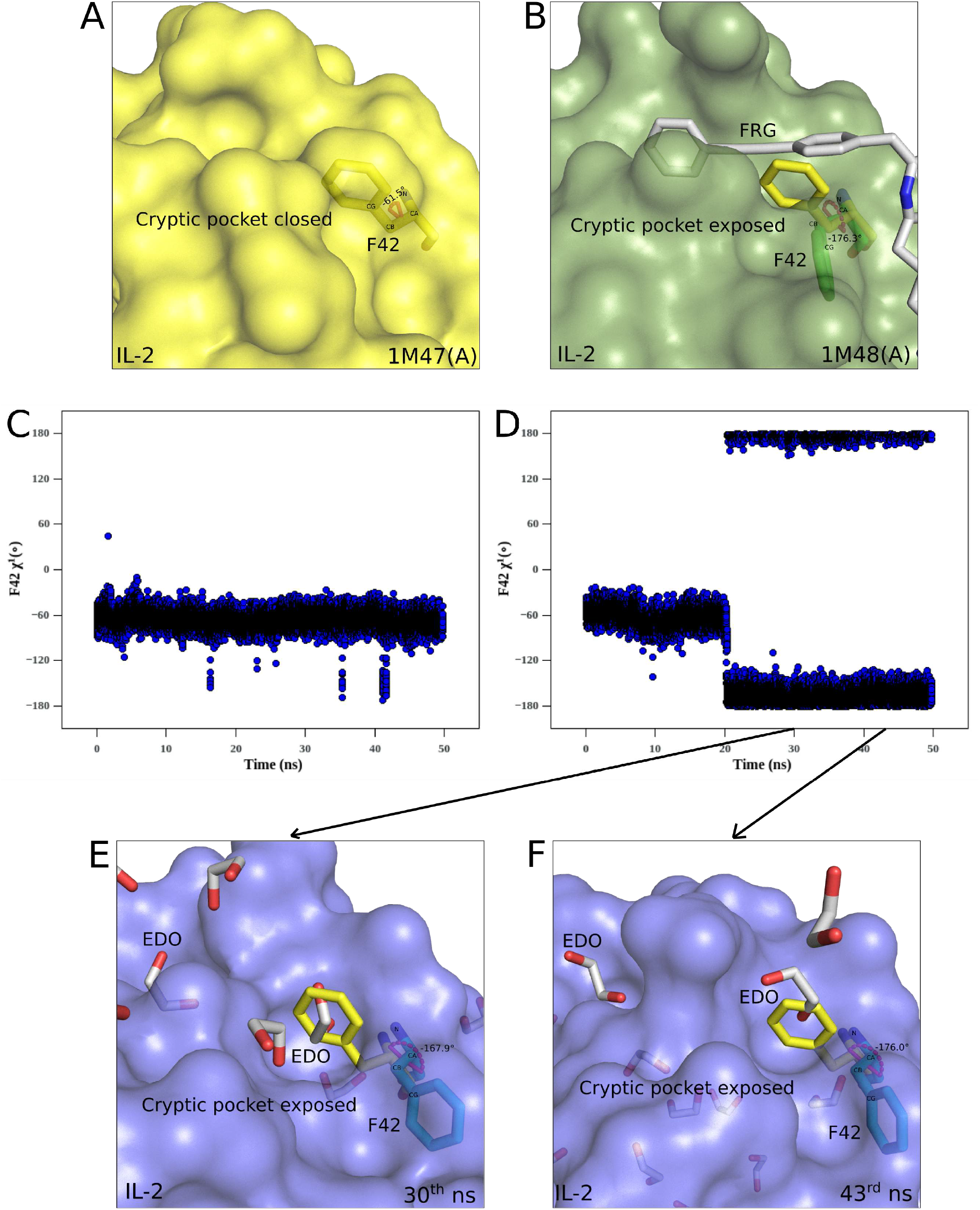
Enhanced sampling of the exposed-state of the cryptic pocket in ethylene glycol based cosolvent simulations of IL-2 / Ethylene glycol based cosolvent simulations enhances sampling of the exposed-state of the cryptic pocket in IL-2. Amino acid residues and bound ligands are shown in a stick model. Nitrogen and oxygen atoms are colored blue and red respectively while carbon atoms are colored according to the color mentioned for residues or ligands. Residue side-chain *χ*^1^ (measured along N,CA,CB,CG atoms) dihedral angles are shown as labeled magenta arcs. Surface of IL-2 is drawn and is superposed with its residue Phe42 to illustrate the conformational change in this residue and the associated opening of the cryptic pocket upon ligand binding. (A) Closed-state of cryptic pocket and associated conformation of residues Phe42 (yellow sticks) is shown in unbound IL-2 structure (yellow, 1M47(A)). (B) Conformational change in residue Phe42 (protruding, yellow sticks; displaced, green sticks) and opening up of the cryptic pocket upon binding of ligand (Ligand ID FRG, grey) to IL-2 structure (smudge, 1M48(C)) is shown. Time series plots showing distribution of the *χ*^1^ side-chain dihedral angle of Phe42, as one of the parameters referring/indicating to the distribution of the closed/open states of the cryptic pocket, in (C) explicit-solvent and (D) 5 % ethylene glycol based explicit-cosolvent MD simulations of apo IL-2 structure (1M47(A)). Snapshots sampled/selected/extracted from the cosolvent simulation trajectory at (E) 30^th^ ns and (F) 43^rd^ ns depicts/shows conformational change in residue Phe42 (protruding, yellow sticks; displaced, blue sticks) and associated opening up of the cryptic pocket upon binding of ethylene glycol molecule(s) (Ligand ID EDO, grey) to IL-2 structure (slate).

Similar to NPC-2, apo-form of the IL-2 structure was subjected to short (50 ns) explicit-solvent and cosolvent MD simulations with ethylene glycol as probe molecules (Methods, Supporting information) to see whether ethylene glycol molecules could enhance sampling of the open-state of the IL-2 cryptic pocket. Comparison of apo and holo structures of IL-2 shows that binding of ligand FRG to IL-2 and opening of the cryptic pocket is accompanied by a significant change in *χ*^1^ (measured along N,CA,CB,CG atoms) dihedral angle of Phe42 from −61.5° in apo IL-2 structure (Figure 6A) to − 176.3° in holo IL-2 structure (Figure 6B). Therefore, transition in Phe42 *χ*^1^ side-chain dihedral angle, as indicator of the dynamics of the cryptic site, was monitored from the explicit-solvent and cosolvent simulation trajectories of IL-2. Time series plots shows that, in comparison to the explicit-solvent simulation of IL-2 (Figure 6C), presence of ethylene glycol molecules in cosolvent simulation significantly/substantially shifts/increases the distribution of Phe42 *χ*^1^ side-chain dihedral angle more towards the values defining the exposed-state of the IL-2 cryptic pocket (Figure 6D). Representative structural snapshots sampled from the cosolvent simulation trajectory at 30^th^ (Figure 6E) and 43^rd^ ns (Figure 6F) depicts/shows that ingression of cosolvent, ethylene glycol molecules, at the cryptic site causes residue, Phe42, to assume conformations (30^th^ ns: Phe42 *χ*^1^ −167.9°, Figure 6E; 43^rd^ ns: Phe42 *χ*^1^ −176.0°, Figure 6F) similar to the site-permissible conformation of this residue (Phe42 *χ*^1^ −176.3°, Figure 6B) observed in the holo-form of the IL-2 structure, thereby unveiling the cryptic pocket (Figure 6E and F). Though the cryptic site of IL-2 opens to a smaller extent in the absence of probe molecules, previous cosolvent simulation study of IL-2 has shown that the 10 % phenol based cosolvent simulation enhanced the sampling of the open-state of the IL-2 cryptic site [31]. Our 5% ethylene glycol based cosolvent simulation of IL-2 shows that the distribution is significantly shifted towards the open-state in comparison to water only simulations of IL-2 (Figure 6C and D). Here, the observation of interest is that a 5% concentration of ethylene glycol is performing on par with a 10 % phenol concentration in cosolvent studies to identify the cryptic pocket of IL-2. Thus, it is seen from our work that ethylene glycol molecules open the cryptic druggable site on pharmaceutical target IL-2.

Though the focus of our cosolvent simulation analysis is to highlight the “cryptic-pocket finding” potential of ethylene glycol molecules, they were also observed to sample physiologically important sites on the proteins and therefore serve as potential candidate for their inclusion as probe molecules in FTmap probe library for computational mapping of ligand-binding hot spots on protiens [17]. We observed that although the non-terminal phenyl moiety of ligand FRG displaces Phe42 and binds to the cryptic pocket (Figure 6B), scenario also observed for ethylene glycol molecules (Figure 6E and F), we also observed in our cosolvent simulations of IL-2 that ethylene glycol molecules were found to bind to a pocket adjacent to the cryptic pocket (Figure 6E and F) which in the holo form of IL-2 is bound by the terminal phenyl moiety of ligand FRG (Figure 6B). This adjacent pocket might serve as an anchor point for the terminal phenyl moiety of ligand FRG enabling the non-terminal phenyl moiety to exploit flexibility at the cryptic site thereby revealing it and pointing to the induced fit mechanism of cryptic pocket identification. Existence of such pockets/sites, termed as hot spots, in the vicinity of cryptic sites, has been suggested as one of the prerequisite for formation of druggable cryptic sites [7] and are exploited by ligands to identify cryptic sites. In fact, a hot spot mapping study on IL-2 has identified this pocket neighboring the cryptic site as one of the strongest hot spot [7]. Thus it can be seen from our cosolvent simulations that ethylene glycol molecules not only samples the cryptic sites but also the neighboring/vicinal hot spots crucial for the identification of druggable cryptic sites.

### “Cryptic-pocket finding” potential of small glycols in other proteins

Further, we analyzed protein structures with validated cryptic sites from Protein Data Bank (PDB) to find out the generality of the “cryptic-pocket finding” potential of these small glycols. Here, we analyze the “cryptic-pocket finding” potential of these glycols in a representative set of diverse proteins, as reported in the Cryptosite set [6, 7] where for each protein from the set, cryptic site was revealed by binding of a biologically relevant ligand to it (Methods, Supporting Information). We found 4 proteins for which, similar to the holo-form of the protein, cryptic site was exposed in the EDO containing structure of that protein and EDO molecules alone were present at that site (Methods, Table S5). Here, we analyze two proteins of these apo-holo protein pairs (rest are discussed in Supporting Information) from the point of view of potential of EDO molecule(s) to bind and occupy the cryptic sites, which were otherwise bound by biologically relevant ligands in the CryptoSite set.

**B-cell lymphoma-extra large (Bcl-xL)** is an anti-apoptotic protein and belongs to the Bcl-2 family of proteins [41]. The binding of BH3 domain of pro-death BH3-only proteins to pro-survival proteins (Bcl-xL), mediated by insertion of four conserved hydrophobic residues (hl-h4) from BH3 domain into hydrophobic pockets (pl-p4) on Bcl-xL surface, initiates apoptosis [42]. ABT-737 (Ligand ID N3C) is a functional BH3 mimetic small-molecule inhibitor of Bcl-2 family proteins, that binds with high affinity to Bcl-xL [43]. In the ABT-737 bound structure of Bcl-xL (2YXJ, chain A) it was observed that the chloro-biphenyl moiety of ABT-737 deepened the p2 pocket of Bcl-xL by displacing side-chains of residues Phe105 and Leu108 thereby further opening-up of the binding groove (Figure 7B), as compared to the p2 pocket in unbound structure of BcL-xL (3FDL, chain A) (Figure 7A). We observed that in the EDO containing structure of Bcl-xL (3FDM, chain A), an EDO molecule also penetrates p2 pocket more deeply and opens-up the binding groove further by displacing side-chains of residues Phe105 and Leu108 (Figure 7C). Furthermore, the comparative analysis of interactions of ABT-737 and EDO with Bcl-xL showed similar hydrophobic contacts of chloro-biphenyl moiety of ABT-737 and the EDO molecule at the enlarged p2 pocket (Figure 7D). Therefore, it may be seen that the EDO molecule, in a manner similar to chloro-biphenyl moiety of ABT-737, also exposes the cryptic site at the p2 pocket of Bcl-xL.

**Figure 7:**
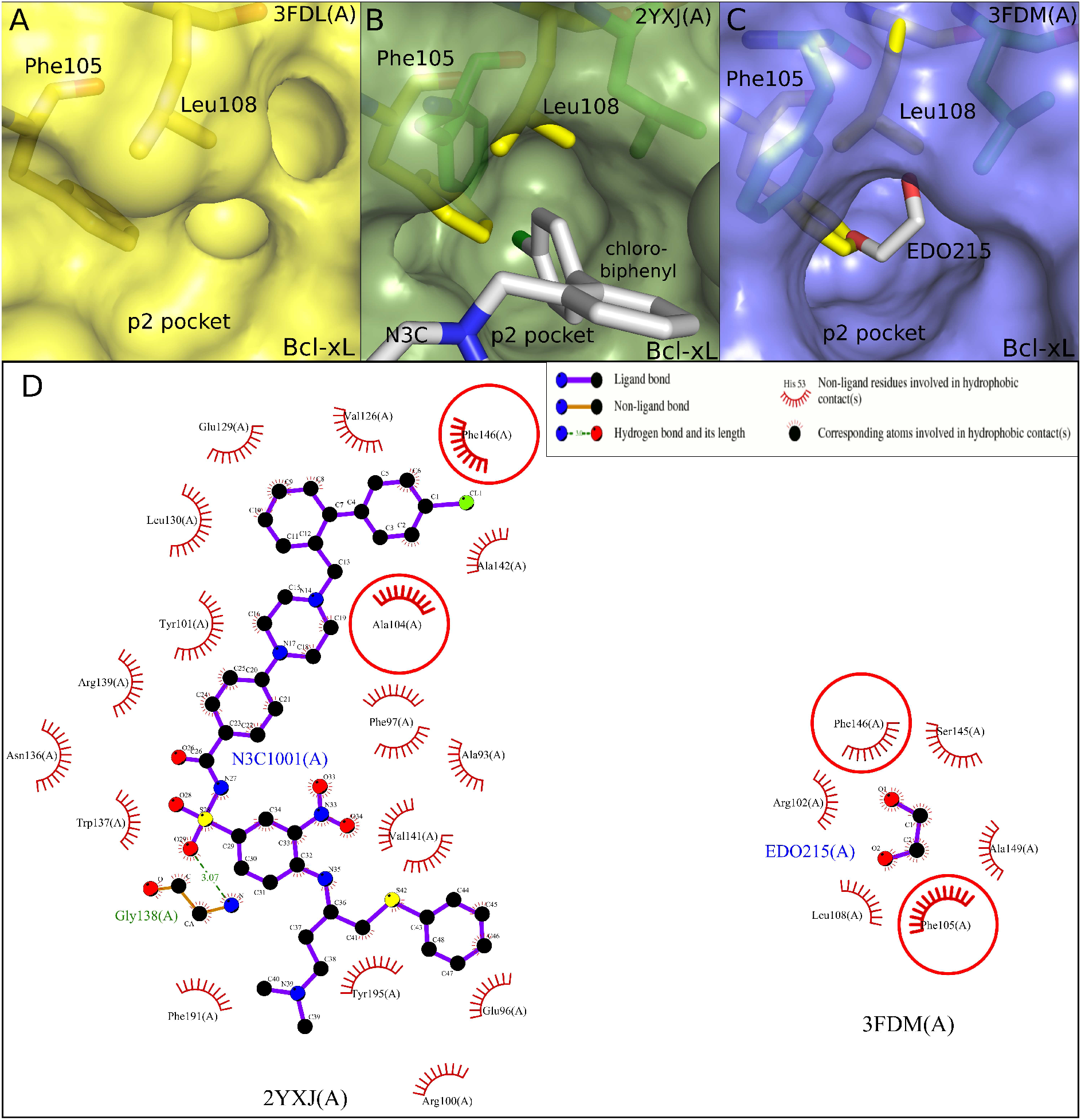
Ethylene glycol identifies a cryptic site in Bcl-xL. Amino acid residues and bound ligands are shown in a stick model. Nitrogen, oxygen and chlorine atoms are colored blue, red and darkgreen respectively while carbon atoms are colored according to the color mentioned for residues or ligands. Surface of Bcl-xL is drawn and is superposed with its residues Phe105, Leu108 to illustrate the conformational changes of these residues and the associated deepening of its p2 pocket upon ligand binding. (A) p2 pocket and conformation of residues Phe105, Leu108 are shown in unbound Bcl-xL structure (yellow, 3FDL(A)). Displacement of protruding side-chains of residues Phe105, Leu108 (protruding, yellow sticks; displaced, green sticks) and enlargement of p2 pocket upon binding of (B) chloro-biphenyl moiety of ligand ABT-737 (Ligand ID N3C, grey) to Bcl-xL (smudge, 2YXJ(A)) and (C) ethylene glycol (Ligand ID EDO, grey) to Bcl-xL (slate, 3FDM(A)) is shown. (D) Schematic representation of the interactions made by ligands ABT-737 (Ligand ID N3C) and ethylene glycol (Ligand ID EDO) with Bcl-xL. Bcl-xL residue identifiers (residue name, number, chain ID) and ligand identifiers (Ligand ID, number, chain ID) along with atom names of bound ligands are indicated. Hydrogen bond length is given in A. The red circles indicate Bcl-xL residues that are in equivalent 3D positions when complexes of Bcl-xL with ligands N3C (2YXJ(A)) and EDO (3FDM(A)) are superposed thereby highlighting the Bcl-xL-N3C and Bcl-xL-EDO interactions that are common at the enlarged p2 pocket of Bcl-xL.

**Myosin II** is a motor protein responsible for producing muscle contraction in muscle cells and plays a central role in other cellular processes such as cell motility and cytokinesis [44]. In the apo-form of Myosin II heavy chain (2AKA, chain A), protrusion of side-chains of residues Leu262 and Tyr634 into the binding pocket occludes it (Figure 8A). Binding of blebbistatin (Ligand Id BIT), a Myosin II inhibitor [45], in the holo-form of the protein (1YV3, chain A) causes these residues to move out of the site thereby exposing it (Figure 8B). In EDO containing structures of Myosin II (1W9J and 1W9L, chain A), the binding of EDO molecule causes side-chains of residues Leu262 and Tyr634 to adopt conformation similar to the site-permissible conformation of these residues observed in the blebbistatin bound structure of Myosin II, thereby identifying the cryptic site (Figure 8C). The binding of blebbistatin to Myosin II involves hydrogen bonding interactions with residues Gly240, Leu262, Ser456 and a number of hydrophobic contacts (Figure 8D). We observed that in EDO containing structures of Myosin II, the binding of EDO molecules at the exposed blebbistatin binding site also involved, among other contacts, hydrogen bonding interactions with either Leu262 or Ser456 or both (Figure 8D). Thus it may be seen that EDO molecules also have the potential to identify the cryptic blebbistatin binding site in Myosin II.

**Figure 8:**
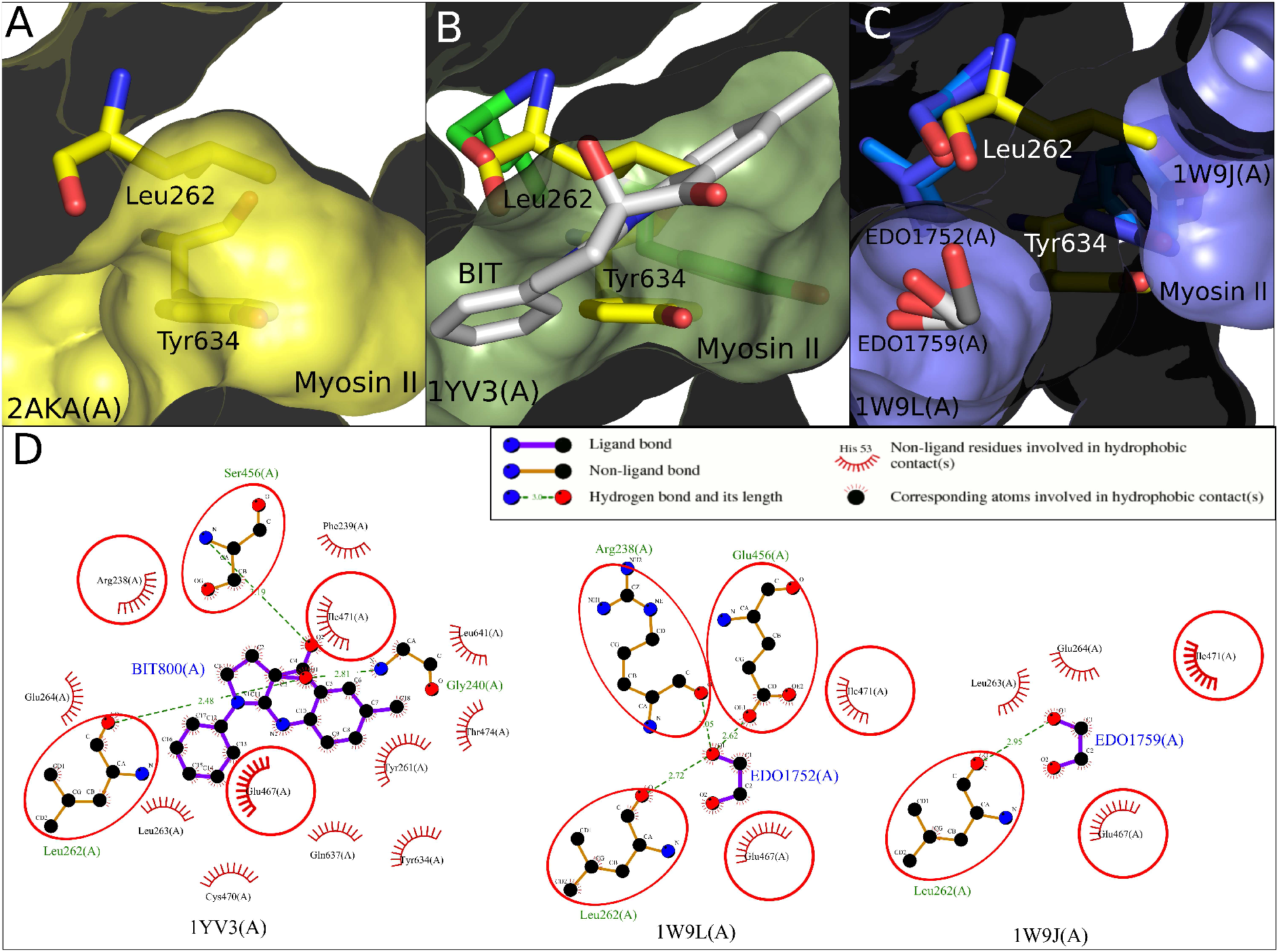
Ethylene glycol identifies a cryptic site in Myosin II. Amino acid residues and bound ligands are shown in a stick model. Nitrogen and oxygen atoms are colored blue and red respectively while carbon atoms are colored according to the color mentioned for residues or ligands. Surface of myosin-II is drawn and is superposed with its residues Leu262, Tyr634 to illustrate the conformational changes of these residues and the associated change in the cryptic site upon ligand binding. (A) Occluded cryptic site due to the protrusion of side-chain of residues Leu262, Tyr634 in unbound myosin-II structure (yellow, 2AKA(A)) is shown. (B) Displacement of protruding side-chains of residues Leu262, Tyr634 (protruding, yellow sticks; displaced, green sticks) and revelation of the cryptic site upon binding of ligand blebbistatin (Ligand ID BIT, grey) to myosin-II (smudge, 1YV3(A)) is shown. (C) Displacement of protruding side-chains of residues Leu262, Tyr634 (protruding, yellow sticks; displaced, blue/darkblue sticks) and revelation of the cryptic site upon binding of ethylene glycol (Ligand ID EDO, grey/darkgrey) to myosin-II (slate, 1W9L(A),1W9J(A)) is shown. (D) Schematic representation of the interactions made by ligands blebbistatin (Ligand ID BIT) and ethylene glycol (Ligand ID EDO), with myosin II. Myosin-II residue identifiers (residue name, number, chain ID) and ligand identifiers (Ligand ID, number, chain ID) along with atom names of bound ligands are indicated. Hydrogen bond length is given in A. The red circles and ellipses indicate myosin-II residues that are in equivalent 3D positions when complexes of myosin-II with ligands BIT (1YV3(A)) and EDO (1W9L(A),1W9J(A)) are superposed thereby highlighting the myosin-II-BIT and myosin-II-EDO interactions that are common at the exposed cryptic site of myosin-II.

It has been shown that in most cases (19.4 %) protrusion of side-chains in the pocket is the cause of pocket being cryptic [7], and for RBSX-W6A, we found that binding of EDO or PGO displaces the site-occluding side-chain of residue Phe4 to reveal the cryptic surface pocket (Figure 1). We also observed that binding of ligands, ABT-737, blebbistatin and ethylene glycol (Ligand IDs N3C, BIT and EDO) to proteins Bcl-xL and Myosin II heavy chain, causes site-occluding side chains of residues Phe105, Leu108 and Leu262, Tyr634 to adopt site-permissible conformations thereby exposing the cryptic site (Figure 7 and 8).

Interestingly, in all the three systems, RBSX-W6A, Bcl-xL and Myosin II heavy chain, EDO molecules are able to displace an aromatic side-chain (Phe4, Phe105 and Tyr634) to identify the cryptic site. For G-actin and Glutamate receptor 2 (discussed in Supporting Information), flexibility at protein carboxy-terminus and loop protrusion in the site are responsible for the sites being cryptic (Figure S5 and S6). Besides the observation that only EDO molecules are present at the exposed cryptic sites in EDO containing structures of G-actin and Glutamate receptor 2 (Figure S5 and S6), other factors may also account for the changes required in carboxy-terminus and loops of these proteins for revealing the cryptic site. Importantly, in all the four protein systems analyzed above, the interaction profile of ethylene glycol shares features with the interaction profile of the ligand at the exposed cryptic site (Figure 7D, 8D, S5D and S6D) thus suggesting that binding at cryptic sites by EDO molecules in these proteins is not by chance but a specific and targeted event. All the four proteins discussed are pharmaceutically important targets. Our analysis thus shows the ability of ethylene glycol molecules to bind to cryptic druggable sites as observed in Bcl-xL and Myosin II proteins (Figure 7 and 8) in addition to the cryptic sites of glutamate receptor 2 and actin.

## Conclusions

Among the effects of polyhydric alcohols on proteins, it has been reported that glycerol and polyethylene glycol interact with features in proteins [46], It is also known that ethylene glycol and propylene glycol bind to proteins and are used as cryo-protectants in protein crystallography experiments. However that small glycols, such as ethylene glycol and propylene glycol, display positive and desirable effect of uncovering cryptic sites in proteins has not been elaborated earlier. The present study has for the first time systematically explored and demonstrated the “cryptie-site finding” potential of these small glycols in proteins. Stand alone small cryptic pockets formed solely by movement of side-chains in general may not be useful for drug design [7], however cryptic adjacent pockets which occur in the vicinity of functional sites of proteins can aid in the design of ligand molecules with increased affinity and specificity, given their identification by some means.

In that respect, small probe molecules such as ethylene glycol and propylene glycol, which in the present work are shown to discover cryptic sites in proteins mainly by inducing side chain movements, can be used to probe the vicinity of functional sites of pharmaceutically important protein targets for potential cryptic sites and/or binding hot spots. Such cryptic sites and/or hot spots, identified vicinal to functional sites, can then be used in a structure-guided/aided chemical elaboration process to design ligands which bind with enhanced affinity to those targets. In fact, upon screening protein CK2*α*, a Ser/Thr kinase and an important target in cancer therapy, with fragment library, a fragment 3,4 dichlorophenethylamine identified a cryptic pocket, termed as “*α*D pocket”, adjacent to ATP binding site by displacing side chain of Tyr125 (PDB ID 5CLP) [47]. This cryptic site was then used to develop a new CK2*α* inhibitor with high nanomolar affinity [47]. Interestingly, in one structure of CK2*α* (PDB ID 3WAR), it was observed that two ethylene glycol molecules were bound at the entrance of the partially opened *α*D pocket [47, 48] further supporting our concept of using small glycols to identify cryptic sites in protein targets which can subsequently be used for drug-design purposes. Another crystallography study of protein CK2*α* [48] has shown several ethylene glycol molecules occupying physiologically significant sites of CK2*α* suggesting their use in computational methods for binding site determination. Thus it must be noted that, for a given protein (in this case CK2*α*), ethylene glycol molecules are able to identify a cryptic site in the protein as well as are able to map the binding hot spots of the protein. Similarly, through crystallography experiments, propylene glycol has been shown to map hot spots of binding in functional sites of various proteins and has been suggested as a relevant seed for further design [49]. Since it is evident from our work that small glycols can unveil cryptic sites, they can be used as fragments in experimental fragment screening methods with added advantage of identifying cryptic sites thereby enhancing the repertoire of mini-fragment libraries. Further, computationally these small glycols can be included in the probe set of hot-spot mapping protocols and mixed-solvent MD simulations to identify cryptic sites and map binding hot-spots on available structures of protein targets as well as on modeled structures of protein targets which are difficult to crystallize. Importantly, our work argues for the use of small glycols in both experimental as well as computational protocols for the identification of cryptic sites which could be eventually exploited in drug-design endeavors.

## Abbreviations

RBSX: Recombinant xylanase from *Bacillus sp. NG-27*
RBSX-W6A: RBSX Tryptophan6 to Alanine mutant
MD: Molecular dynamics
PDB: Protein Data Bank
EDO: 1,2-ethanediol (ethylene glycol)
PGO: S-1,2-propanediol (propylene glycol)
PGR: R-1,2-propanediol (propylene glycol)
AwSA: atom-wise solvent accessibilities
Bcl-xL: B-cell lymphoma-extra large
TMR: tetramethylrhodamine-5-maleimide
ITC: isothermal titration calorimetry

## Acknowledgements

The authors thank CSIR-UGC and the Indian Institute of Science (IISc) for providing financial support to H.B., UGC for emeritus fellowship to S.R., X-ray Facility at the Molecular Biophysics Unit (IISc), Bengaluru, India and European Synchrotron Radiation Facility (ESRF), Grenoble, France for X-ray data collection, staff for providing access to and support on the beamline BM14 at ESRF, DBT for funding trip to ESRF, DST sponsored computational facility in the Department of Physics (IISc) and Supercomputer Education and Research Center (IISc) for providing access to CRAY XC_40_-“SAHASRAT” supercomputer and other computational resources for MD simulations and Dr. V.S. Reddy and Dr. Amit Bharadwaj for providing protein samples of RBSX and RBSX-W6A. The authors thank Ms. Julia Fecko for the ITC binding study. The ITC work was supported by the NIH grant S100D025145 to Dr. Yennawar for the TA Instruments Low Volume AutoAffinity ITC, housed in the Automated Biological calorimetry core facility at the Penn State, Huck Institutes of the Life Sciences.

## Author Contributions

H.B., P.M. and S.R. conceptualized the study. H.B. and P.M. crystallized and solved the crystal structures. H.B. carried out MD simulations and other computational studies. N.H.Y. performed ITC experiments. H.B., P.M., N.H.Y., and S.R. analyzed and interpreted the data. H.B. wrote the paper which was reviewed, edited and approved by all the coauthors. In the opinion of all authors, H.B. and P.M. should be considered as first authors.

## Notes

**Conflicts of interest:** There are no conflicts of interest to declare.

### Competing Interest Statement

The authors have declared no competing interest.

