## Supplementary material for "Small Glycols Discover Cryptic Pockets on Proteins for Fragment-based Approaches": Table S1, Table S2, Table S3, Table S4, Table S5, Figure S1, Figure S2, Figure S3, Figure S4, Figure S5, Figure S6

### Methods

#### Crystallization and data collection

The purification of RBSX-W6A was carried out as described previously as a part of project investigating the stability of proteins [1]. RBSX-W6A was crystallized in sitting drop vapor diffusion plates at a protein concentration of ~10 mgml$^{-1}$ by mixing 1 µl of protein solution with 1 µl of mother liquor. The plates were kept at 20 °C-22 °C. The condition, 100 mM NaCL, 160 mM $MgCl_2$, 50 mM Tris-HCl pH 8.5 and 20 % PEG 8000, gave RBSX-W6A crystals in a week. Prior to data collection, the RBSX-W6A crystals were first soaked for 2-3 minutes in mother liquor solution containing 10 % (v/v) ethylene glycol (Ligand ID EDO) or 10 % (v/v) propylene glycol and subsequently flash-cooled in liquid nitrogen. Ethylene glycol was used as a cryo-protectant. Synchrotron data set for RBSX-W6A crystal soaked with ethylene glycol was collected on a MarCCD detector at the European Synchrotron Radiation Facility (ESRF) beamline BM14, at 100 K with a wavelength of 0.82656 Å. Home source data sets for RBSX-W6A crystal and RBSX-W6A crystal soaked with propylene glycol were collected on a mar345 image-plate detector system at X-ray Facility for Structural Biology, Molecular Biophysics Unit, Indian Institute of Science, at 100K with a wavelength of 1.5418 Å.

#### X-ray data processing, structure determination and refinement

Indexing and integration of all the X-ray data sets was carried out using iMOSFLM [2] followed by scaling and merging with AIMLESS [3] program in CCP4 program suite [4]. Molecular replacement using the program Phaser-MR [5] from the CCP4 program suite was used for structure solution with native RBSX structure (PDB code 4QCE) [6] as the search model. Automated model rebuilding and completion of the molecular replacement solution was carried out using Phenix AutoBuild Wizard which uses RESOLVE, xtriage and phenix.refine to build an atomic model, refine it, and improve the same with iterative density modification, refinement, and model building [7]. Further rounds of manual model building and refinement were carried out using COOT[8] and REFMAC [9, 10] from the CCP4 program suite. For cross-validation, 5 % of randomly selected reflections were kept aside and were not used in refinement. Final models were validated using MolProbity [11] and their stereochemistry was analyzed with PROCHECK [12].

Table S1: **Statistics of X-ray diffraction data collection and structure refinement.**

| **PDB Code** | **5EFD**[a] | **5XC0**[a] | **5XC1**[a] |
|---|---|---|---|
| Temperature (K) | 100 | 100 | 100 |
| Wavelength (Å) | 0.82656 | 1.5418 | 1.5418 |
| Resolution range (Å) | 32.10-1.67(1.76-1.67) | 27.96-2.32(2.4-2.32) | 30.01-2.26(2.33-2.26) |
| Space group | $P2_12_12_1$ | $P2_12_12_1$ | $P2_12_12_1$ |
| Unit Cell Dimensions | | | |
| $a$, $b$, $c$ (Å) | 54.99,76.60,176.49 | 52.23,67.39,177.76 | 55.0,75.73,176.60 |
| $\alpha$, $\beta$, $\gamma$ (°) | 90,90,90 | 90,90,90 | 90,90,90 |
| Unit cell volume (Å$^3$) | 743417 | 625667 | 735558 |
| Solvent content (%) | 45.10 | 35.63 | 45.25 |
| Unique reflections | 86543 | 27309 | 35296 |
| $R_{merge}$[b] (%) | 7.7(42.1) | 14(45.2) | 12.1(44.5) |
| No. of molecules in asymmetric unit | 2 | 2 | 2 |
| $V_M$(Å$^3$Da$^{-1}$) | 2.24 | 1.91 | 2.25 |
| Wilson B-factor (Å$^2$) | 14.5 | 20.0 | 20.6 |
| Multiplicity[b] | 5.7(5.3) | 5.3(5.5) | 6.9(6.3) |
| Average I/$\sigma$I[b] | 14.7(4.0) | 9.2(3.6) | 11.7(3.5) |
| Completeness (%)[b] | 99.8(99.3) | 97.7(93.0) | 99.7(97.2) |
| $R_{work}$/$R_{free}$ | 15.68/18.58 | 20.2/24.8 | 18.8/22.7 |
| No. of atoms | | | |
| Protein | 5842 | 5710 | 5764 |
| Ligand/ion | 33 | 4 | 14 |
| Water | 442 | 209 | 393 |
| Root mean square bond lengths (Å) | 0.0139 | 0.0116 | 0.0163 |
| Root mean square bond angles (°) | 1.5995 | 1.3982 | 1.6022 |
| **Average B-factors Å$^2$** | | | |
| Protein | 16.7 | 27.2 | 22.5 |
| Ligand/ion | 21.4 | 9.6 | 31.6 |
| Water | 25.9 | 17.5 | 24.2 |
| **Ramachandran map statistics (%)** | | | |
| Most favored regions | 91.7 | 91.9 | 92.7 |
| Additional allowed regions | 8.3 | 8.1 | 7.3 |
| Generously allowed regions | 0 | 0 | 0 |
| Disallowed regions | 0 | 0 | 0 |
| [a]**Cryptic pocket status** | open containing ethylene glycol | closed | open containing propylene glycol |

[b]Highest resolution shell is shown in parentheses.

Detailed statistics of data collection, refinement and Ramachandran plot are presented in Table S1. The atomic coordinates and structure factors of all the three structures have been deposited by us in Protein Data Bank (PDB) and are available under the accession codes 5EFD, 5XC0 and 5XC1.

#### Analysis of structures

The pocket lining atoms and their atom-wise solvent accessibilities (AwSA) were calculated using the Pocket program, part of ProShape software (http://biogeometry.cs.duke.edu/software/proshape/software.html) and the NACCESS program respectively [13] (Table S2). Contacts were found using NCONT program from the CCP4 program suite [4]. Schematic drawings of protein-ligand interactions were generated using LigPlot+ [14]. Superposition of structures and generation of molecular graphics were done using PyMOL [15].

Table S2: **Atoms lining the cryptic surface pocket and their their atom-wise solvent accessibilities (AwSA) in 5EFD (Chain A), 5XC0 (Chain B) and 5XC1 (Chain B).**

| **Residue** | **5EFD (Chain A)**<br>**Atoms : AwSA (Å$^2$)** | **5XC0 (Chain B)**<br>**Atoms : AwSA (Å$^2$)** | **5XC1 (Chain B)**<br>**Atoms : AwSA (Å$^2$)** |
|---|---|---|---|
| PHE4 | CB : 7.995<br><br>CD1 : 4.298 | CB : 13.636<br>CG : 1.729<br>CD1 : 4.156 | CB : 9.857<br>CG : 2.171<br>CD1 : 5.127 |
| ALA5 | N : 0.633<br>CB : 0.045 | N : 0.000<br>CB : 0.097 | N : 0.420<br>CB : 0.238 |
| **ALA6** | **N : 1.059**<br>CB : 14.814 | **N : 0.000**<br>CB : 5.433 | **N : 1.393**<br>CB : 15.584 |
| ARG33 | CA : 6.333<br>CB : 26.250<br>O : 0.016 | CA : 2.932<br>CB : 23.458<br>O : 0.000 | CA : 3.145<br>CB : 35.902<br>O : 0.296 |
| LYS36 | CB : 6.567<br>CG : 11.055<br>CD : 28.699 | CB : 6.072<br>CG : 9.777<br>CD : 18.549<br>NZ : 47.153 | CB : 7.209<br>CG : 11.672<br>CD : 27.344<br>NZ : 44.090 |
| VAL37 | CG2 : 3.869 | CG2 : 0.007 | CG2 : 4.338 |
| TYR343 | CB : 5.030<br>CD2 : 6.077 | CB : 0.262<br>CD2 : 6.300 | CB : 6.195<br>CD2 : 8.048 |
| **Cryptic pocket status** | open containing ethylene glycol | closed | open containing propylene glycol |

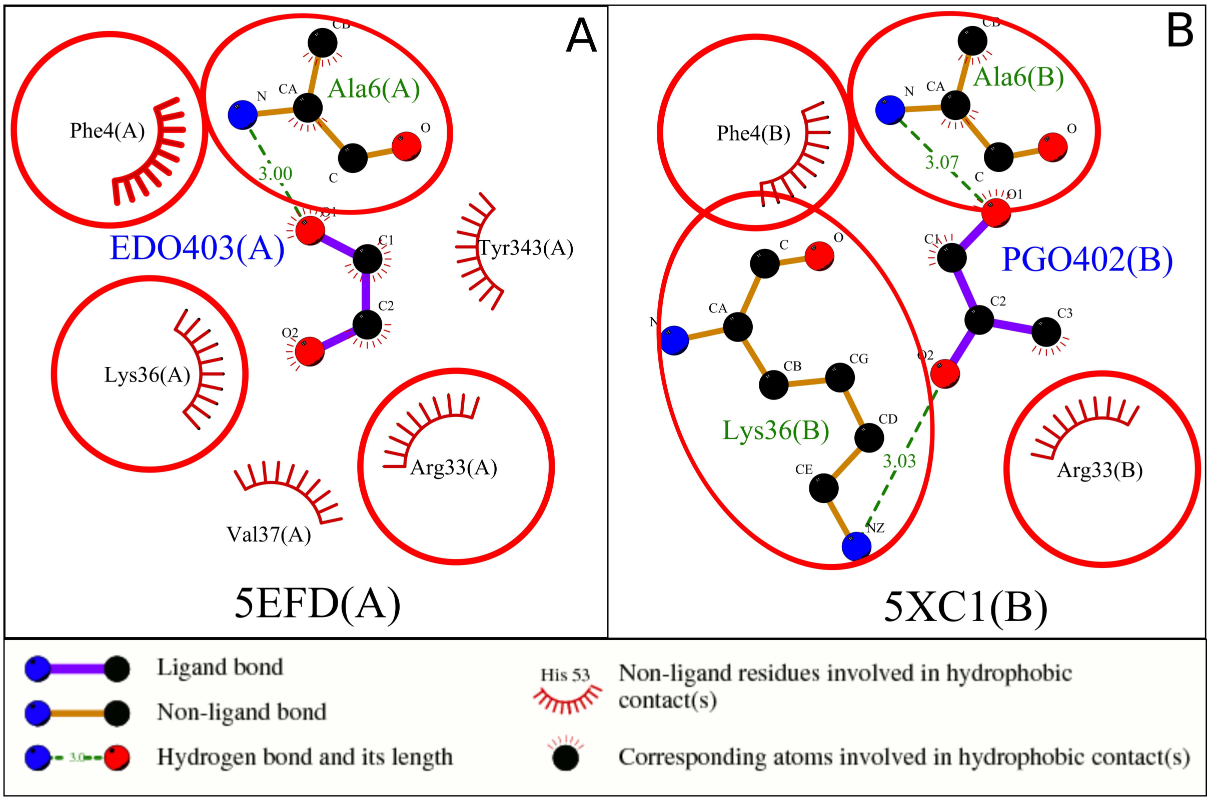


Figure S1: **Schematic representation of the interactions made by different small glycols at the exposed cryptic surface pocket of RBSX-W6A.** (A) Ethylene glycol (EDO) and RBSX-W6A (5EFD(A)) (B) Propylene glycol (PGO) and RBSX-W6A (5XC1(B)). RBSX-W6A residue identifiers (residue name, number, chain ID) and glycol identifiers (Ligand ID, number, chain ID) along with atom names of bound glycols are indicated. Hydrogen bond length is given in Å. The red circles and ellipses indicate RBSX-W6A residues that are in equivalent 3D positions when complexes of RBSX-W6A with glycols EDO (5EFD(A)) and PGO (5XC1(B)) are superposed thereby highlighting the RBSX-W6A-EDO and RBSX-W6A-PGO interactions that are common at the exposed cryptic surface pocket of RBSX-W6A.

#### Isothermal Titration Calorimetry (ITC)

ITC binding study was performed at 25 °C using standard ITC procedures [16] using the TA instruments AffinityITC Auto equipment. Prior to ITC, lyophilized RBSX and RBSX-W6A proteins and ethylene glycol were dissolved in 25 mM Tris, pH 8.5, to a final concentration of 160 µM. In order to obtain a good reference run, the protein was used as the titrant in the syringe with ethylene glycol in the calorimeter cell. For each sample ITC run, 2 µl of the protein were injected into 160 µM ethylene glycol at 180 second intervals while the contents of the cell were stirred at a speed of 125 rpm.

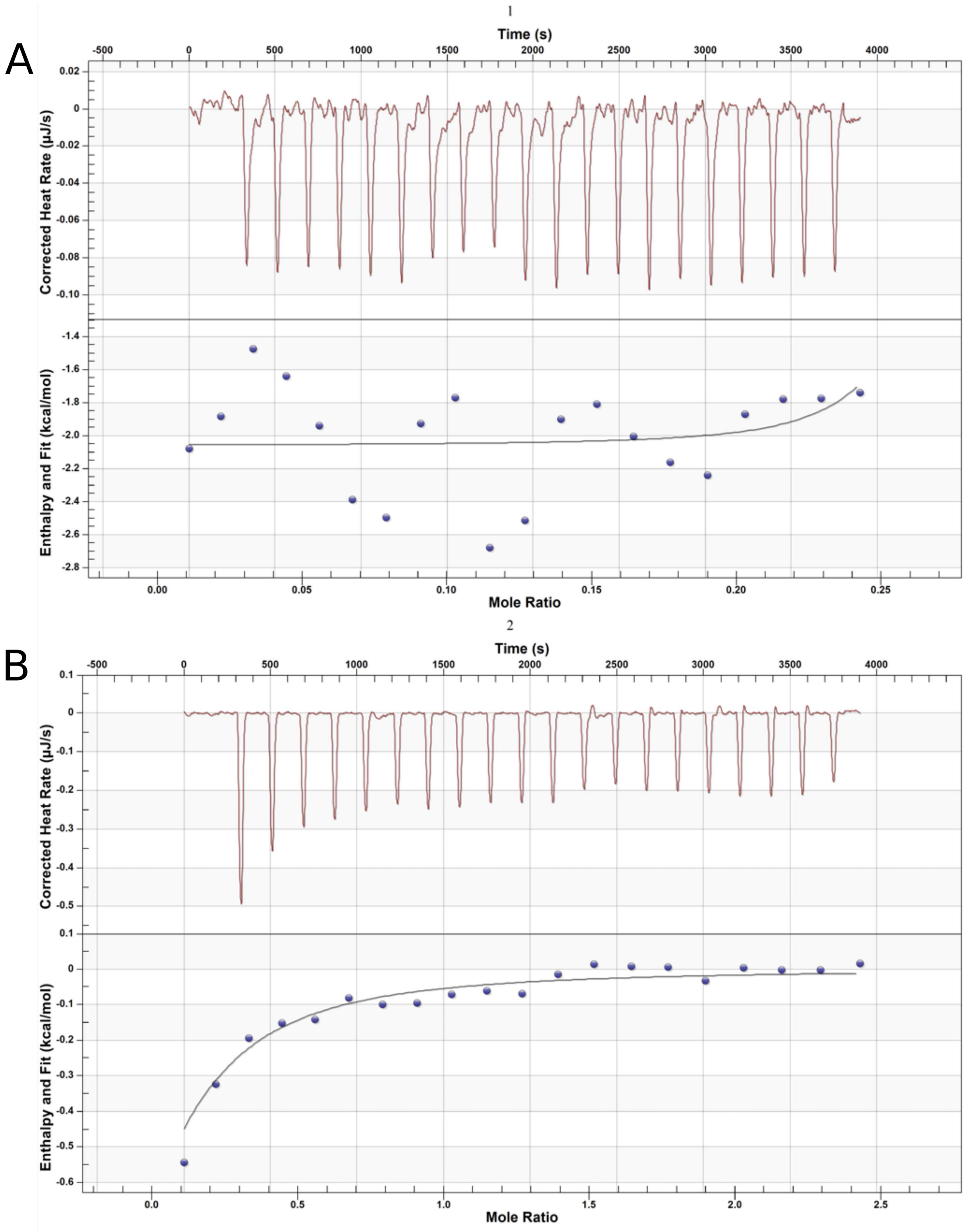


Figure S2: **Isothermal Titration Calorimetry (ITC) monitors the thermodynamics of (A) RBSX and (B) RBSX-W6A binding interactions with ethylene glycol.** Representative power-response curves (top) and heats of reaction normalized to the moles of protein injected (bottom) are provided for each run. All titrations were conducted at 298K in 25 mM Tris, pH 8.5 .

For the reference subtraction, just the buffer was injected into 160 µM ethylene glycol. The best-fit parameters from a single-site binding model are reported with their associated uncertainties represented by the standard error of the mean (Figure S2, Table S3). All data was fit in the Nanoanalyze software (TA instruments).

Table S3: **Thermodynamic parameters describing the binding of RBSX and RBSX-W6A to ethylene glycol measured at 298K in 25 mM Tris, pH 8.5.** All error bars on fitted ITC parameters represent standard error of the mean estimated with fitting performed in the Nanoanalyze software (TA instruments). No measurable binding was seen between ethylene glycol and the RBSX while weak binding was detected for the RBSX-W6A.

| **Ethylene glycol** | **N (sites)** | **Kd (M)** | **ΔH (kcal/mol)** | **ΔS (cal/mol*K)** | **ΔG (kcal/mol)** |
|---|---|---|---|---|---|
| **RBSX** | Not detected | Not detected | Not detected | Not detected | Not detected |
| **RBSX-W6A** | 0.10 (0.16) | 5.70 e-5 (2.20 e-5) | -2.47 (1.18) | 11.14 | -5.79 |

#### System preparation for explicit-solvent and cosolvent molecular dynamics simulations

The structures of RBSX-W6A with the cryptic site in occluded-state (PDB ID 5XC0, chain B) and exposed-state (PDB ID 5EFD, chain A, EDO removed), associated with the closed and open state of residue Phe4, were used as starting structures for explicit-solvent molecular dynamics (MD) simulations. The apo structures of Niemann-Pick type C2 (NPC-2) (PDB ID 1NEP) and interleukin-2 (IL-2) (PDB ID 1M47) were used as starting structures for explicit-solvent and cosolvent MD simulations. All the structures were retrieved from the PDB and files were prepared by first removing all the solvent molecules including any bound ligands. Missing atoms and residues (IL-2) were modelled using Modeller [17]. For explicit-solvent MD simulations, the cleaned structures were solvated in a cubic TIP3P water box with a padding of 10 Å from all sides and neutralized by adding required number of counter ions using the TLEAP module of AmberTools16 [18]. NPC-2 and IL-2 proteins were subjected to explicit-cosolvent MD simulations using ethylene glycol as a probe molecule.

Projects W-26, W-29 and W-30 corresponding to trans, gauche$^-$ and gauche$^+$ conformations of ethylene glycol molecules were used from the R.E.D. server database [19] to obtain the RESP (restrained electrostatic potential) atomic charges for ethylene glycol. Antechamber from Amber 14 software package [20] was then used to derive atom type definition and the associated force field parameters of ethylene glycol using the GAFF force field [21]. For cosolvent MD simulations, NPC-2/IL-2 were solvated in a box containing 5 % v/v cosolvent of ethylene glycol-water molecules and neutralized by adding required number of counter ions using the TLEAP module of AmberTools16 [18]. Ratios of number of ethylene glycol molecules to water molecules used for 5 % v/v probe-water solution are given in the Table S4. First, a shell of probe molecules corresponding to required number of ethylene glycol molecules was placed around the NPC-2/IL-2 protein (Figure S3A and C) using the program Packmol [22]. NPC-2/IL-2 protein coated with probe molecules were then placed in TIP3P box containing water molecules (Figure S3B and D). Volume of the box and thereby the number of water molecules was adjusted to obtain the 5 % v/v ethylene glycol-water cosolvent.

Table S4: **Ratio of ethylene glycol and water molecule in 5 % v/v ethylene glycol-water solutions for cosolvent simulations of NPC-2 and IL-2 proteins.**

| | **Ethylene glycol[1] : Water[2]** |
|---|---|
| **Volume : Volume** | 5 : 95 |
| **Weight : Weight** | 5.55 : 95 |
| **Mol : Mol** | 1 : 59 |
| **Molecule : Molecule** | 105 : 6195 |

[1]Ethylene glycol: 62.07 g/mol;
[2]Water: 18.02 g/mol

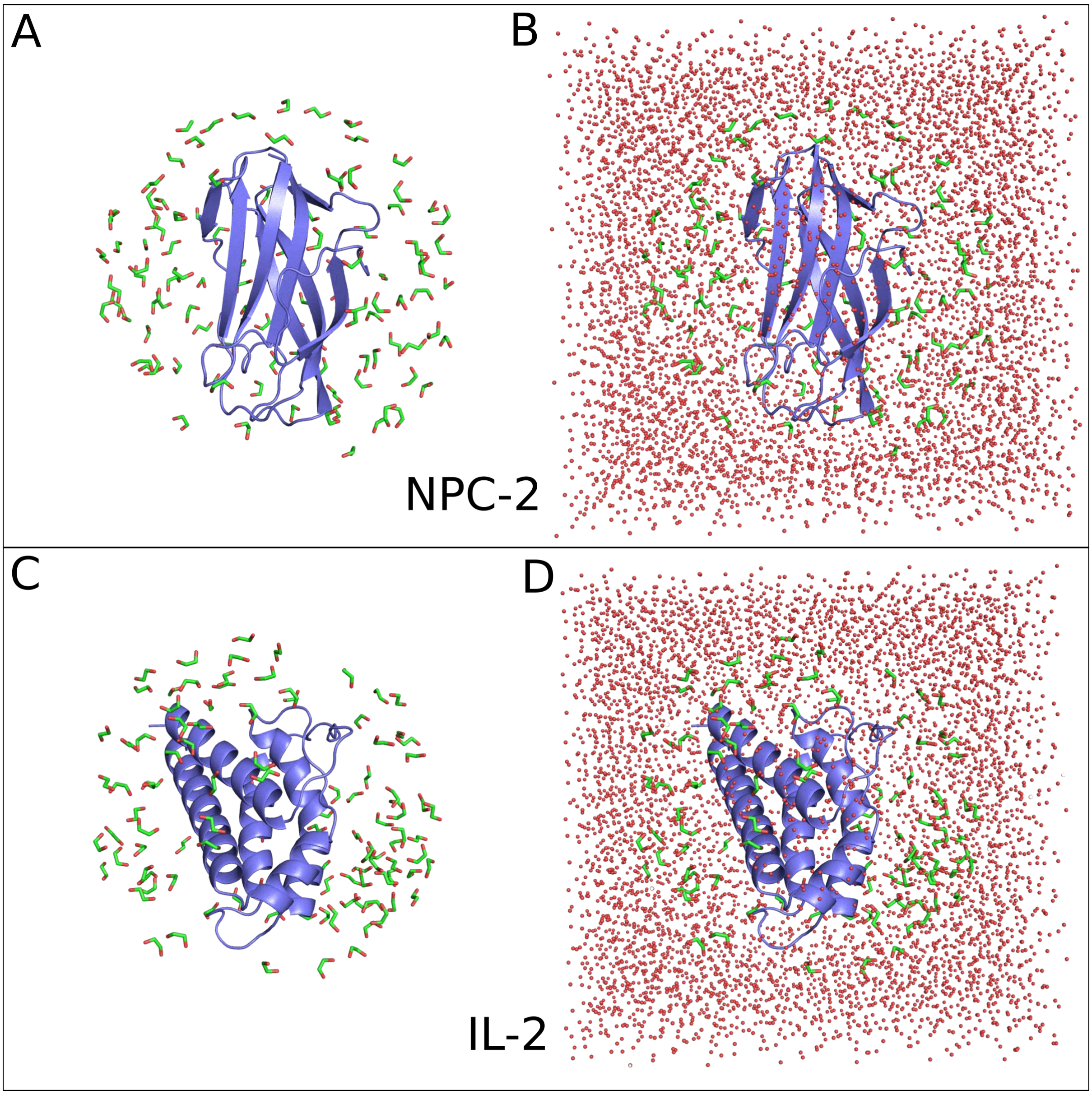


Figure S3: **System setup for ethylene glycol based explicit-cosolvent MD simulations.** Cartoon for protein (slate), sticks for ethylene glycol (EDO) molecules (green) and spheres for water molecules (red) is shown. (A) NPC-2 with EDO (B) NPC-2 with EDO and water (C) IL-2 with EDO (D) IL-2 with EDO and water.

#### Molecular dynamics simulations

All the MD simulations were carried out with the PMEMD module in the Amber 14 software package [20] using the ff14SB force field [23]. Three series of energy minimization were carried out on solvated systems using the steepest descent (5000 steps) and conjugate gradient (5000 steps) algorithms with positional restraints of 100, 50 and 5 kcal/mol.Å$^2$ applied to all solute atoms. This was followed by unrestrained minimization without any positional restraints. The minimized systems were then gradually heated to a temperature of 300 K in 200 ps with positional restraint of 5 kcal/mol.Å$^2$ applied to all solute atoms using the $NVT$ (constant number, volume, and temperature) ensemble. Subsequently the systems were equilibrated for a period of 1 ns and subjected to the production MD runs using the $NPT$ (constant number, pressure (1 atm), and temperature (300 K)) ensemble and periodic boundary conditions. Langevin dynamics [24, 25] with a collision frequency of 2.0 ps$^{-1}$ and Berendsen barostat [26] with pressure relaxation time of 1 ps were used to control temperature and pressure respectively. Particle Mesh Ewald method [27] was used to treat long-range interactions with a 9 Å nonbonded cutoff. SHAKE algorithm [28] was applied to constrain all bonds involving hydrogens, with an integration time step of 2 fs.

#### Analysis of simulation trajectories

CPPTRAJ trajectory analysis program [29] was used for calculating the radial distribution function of water, the side-chain $\chi^1$, $\chi^2$ dihedral angles and atom-atom distance of residues involved in opening and closing of cryptic sites from the explicit-solvent and cosolvent simulation trajectories of RBSX-W6A, NPC-2 and IL-2 proteins. Superposition of structures and generation of molecular graphics were done using PyMOL [15]. The backbone root mean square deviation (RMSD) plots for trajectories of all simulated systems are given in Figure S4. The plots depicts that all simulated systems are well behaved during simulations with minimal RMSD fluctuations.

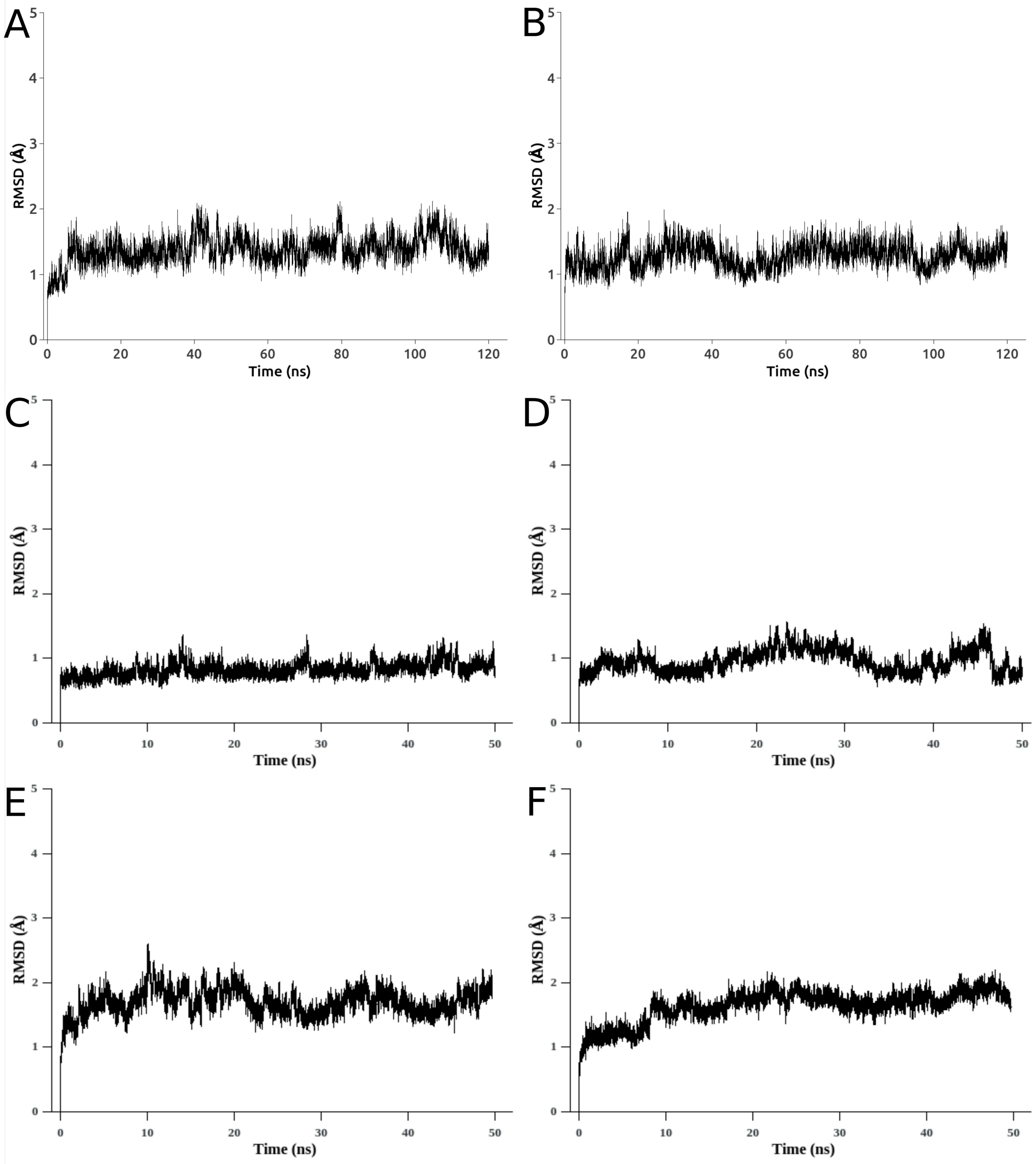


Figure S4: **Backbone RMSD to the starting structure of** (A) apo RBSX-W6A (B) holo RBSX-W6A (C) apo NPC-2 (D) apo NPC-2 with EDO molecules (E) apo IL-2 (F) apo IL-2 with EDO molecules, over varied explicit-solvent and cosolvent MD simulations time scales. Here, apo/holo refers to the unbound/bound and therefore closed/open state of the respective cryptic pockets.

#### "Cryptic-pocket finding" potential of small glycols in other proteins: Dataset construction and analysis

We constructed a dataset of proteins to investigate the "cryptic-pocket finding" potential of small glycols in other proteins. CryptoSite set [30, 31] is a dataset of 93 pairs of apo-holo protein X-ray structures with validated cryptic binding sites such that presence of a biologically relevant ligand in the holo-form of the protein reveals a cryptic site not apparent in the apo-form of that protein. CryptoSite set was used to retrieve PDB identifiers of apo-holo protein pairs and the Ligand IDs of corresponding biologically relevant ligands responsible for revealing the cryptic sites in the holo-form of the proteins. For the analysis, we first identified apo-holo protein pairs from the CryptoSite set, such that the protein of the identified apo-holo protein pair has additional structures in the PDB containing atleast one of the glycol molecule (ethylene glycol/propylene glycol). For a given protein of apo-holo pair in CryptoSite set, its corresponding UniProt ID was used to list all the structures available for that protein in the PDB. The PDB identifiers of structures containing atleast one of the glycol molecule (ethylene glycol/propylene glycol) were then selected from the above generated list and added to the PDB identifiers of apo-holo pair for that protein in the CryptoSite set. This process was repeated for all the 93 apo-holo protein pairs in the CryptoSite set resulting in 29 apo-holo protein pairs (data not shown) for which each protein from the pair had additional structures in the PDB containing ethylene glycol molecules. However none of the protein of apo-holo protein pair from CryptoSite set was found to have additional structures in the PDB containing propylene glycol molecules. It must be noted that, during the construction of CryptoSite set, solvent molecules, buffer components and crystallographic additives were not considered as ligands [30] and therefore EDO containing structures added to the list above, are not part of the CryptoSite set. Next, for each protein from the list of 29 apo-holo protein pairs identified above, we structurally superimposed its EDO containing structure with the structures of its corresponding apo-holo protein pair. Then, the status of the cryptic site in the EDO containing structure was examined to find out whether EDO molecules are present at the cryptic site, which not apparent in the apo-form of that protein is revealed in the holo-form of that protein upon binding of a biologically relevant ligand.

The resulting apo-holo and EDO containing structures, such that the cryptic site revealed upon binding of a biologically relevant ligand in holo-form of the protein and also bound by EDO molecules in the EDO containing structure(s) of that protein are listed in Table S5 and were used for further analysis.

Table S5: **The PDB identifiers of protein systems in unbound, ligand-bound, EDO-bound forms and corresponding Ligand-IDs where ligand and EDO molecules identify the same cryptic site not apparent in unbound structure.**

| Protein system | Unbound | Ligand bound | Ligand ID | EDO bound |
|---|---|---|---|---|
| **Bcl-xL** | 3FDL(A) | 2YXJ(A) | N3C | 3FDM(A) |
| **Actin** | 1RDW(X) | 1J6Z(A) | RHO | 4B1V(A), 4B1V(B) |
| **Myosin II** | 2AKA(A) | 1YV3(A) | BIT | 1W9J(A), 1W9L(A) |
| **Glutamate receptor 2** | 1MY0(B), 1MY1(C) | 1N0T(D), 1FTL(A) | AT1, DNQ | 4H8J(B), 4H8J(D) |

**Actin** is a protein that can transition between monomeric (G-actin) and filamentous (F-actin) states and plays a critical role in many cellular functions including cell motility, maintenance of cell shape and polarity and transcription regulation [32]. In the apo-form of actin (1RDW, chain X), residues 373-375 at the carboxy-terminus of the protein protrude into a pocket occluding it (Figure S5A). However, in the holo-form of actin (1J6Z, chain A) a fluorescent probe tetramethylrhodamine-5-maleimide (TMR) (Ligand Id RHO) is bound at the exposed pocket although these carboxy-terminus residues were not observed due to disorder (Figure S5B). TMR, bound at the exposed pocket, forms hydrogen bonds with actin residues Tyr133 and Tyr143 and a number of hydrophobic contacts with other residues (Figure S5D). Surprisingly, in the EDO containing structure of G-actin (4B1V), we observed that despite the presence of residues 373-375 at its carboxy-terminus, the cryptic pocket was in exposed-state in 4B1V (chains A and B) and an EDO molecule was present at that pocket indicating its potential to identify the cryptic site (Figure S5C). Furthermore, similarly to TMR, EDO molecule was also observed to interact with the pocket through hydrogen bonds with actin residues Tyr133 and Tyr143 (Figure S5D) showing its affinity with the cryptic site.

Moreover, we observed that in the EDO containing structure of G-actin (4B1V), despite the presence of other polyhydric alcohol such as trihydric alcohol glycerol, it is the molecule of dihydric alcohol ethylene glycol that occupied and interacted with the exposed cryptic site further emphasizing the inherent potential of EDO molecules to identify and interact with cryptic sites.

**Glutamate receptor 2** is a ligand-gated ion channel in the mammalian nervous system and plays an important role in excitatory synaptic transmission [33]. In the apo-form of Glutamate receptor 2 (1MY0, chain B and 1MY1 chain C) hinge motion of domains causes loops (59-67,136-142 and 68-73,139-143) to protrude in the binding site occluding it (Figure S6A) and upon binding of antagonists, ATPO and DNQX (Ligand Id: AT1 and DNQ), the loops move back in the holo-form of the protein (1N0T, chain D and 1FTL, chain A) exposing the cryptic site [31] (Figure S6B). We found that, in EDO containing structure (4H8J) of Glutamate receptor 2, conformation of the site-occluding loops was similar to the site-permissible conformation observed in the holo-form and EDO was the sole molecule occupying the exposed cryptic site in 4H8J (chains B and D) (Figure S6C). The comparative analysis of interactions of ATPO, DNQX and EDO with Glutamate receptor 2 revealed that ATPO, DNQX and EDO engages common Glutamate receptor 2 residues upon binding to the exposed site (Figure S6D). Thus, EDO molecules are able to bind and interact with the cryptic site.

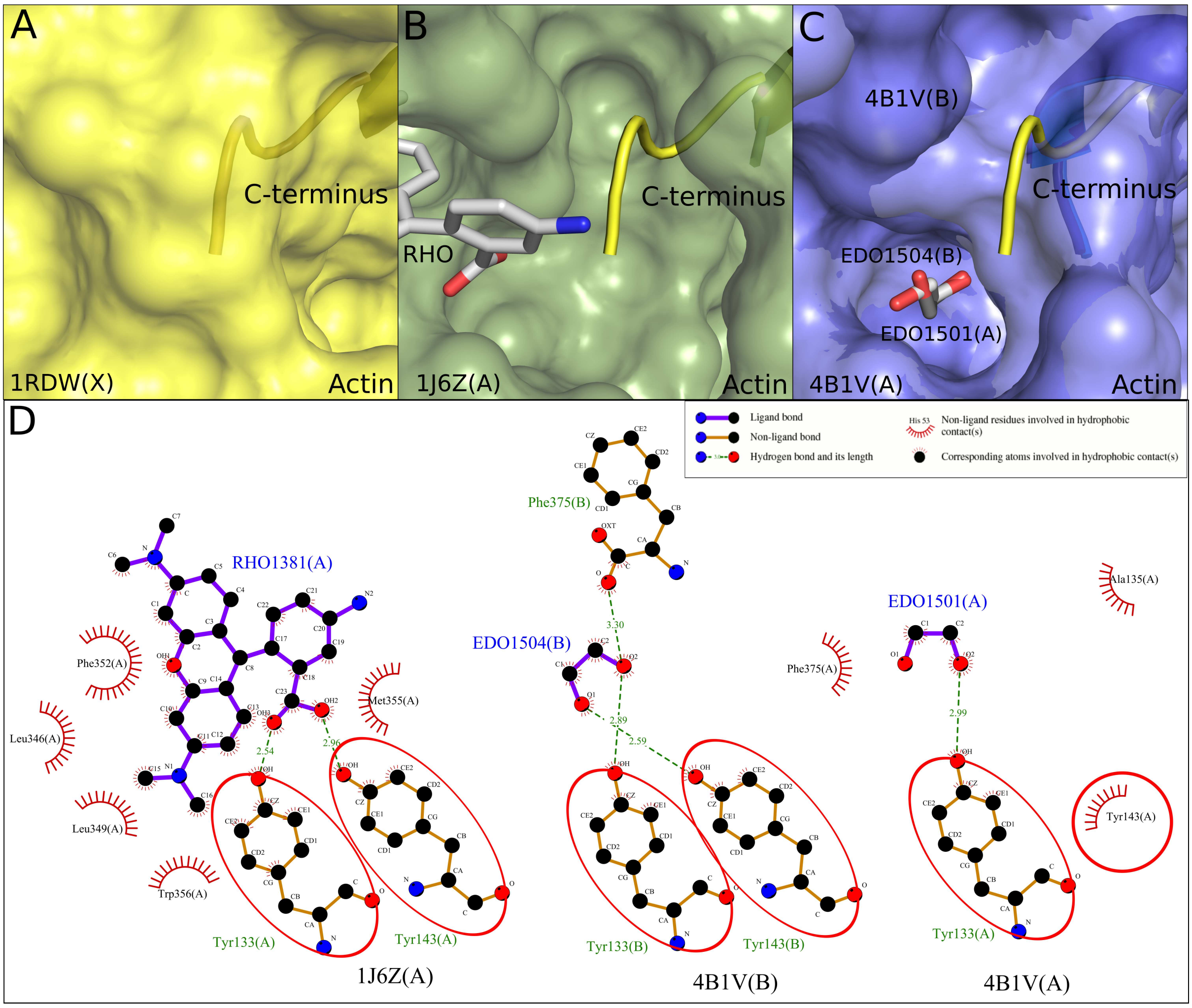


Figure S5: **Ethylene glycol identifies a cryptic site in Actin.** Bound ligands are shown in a stick model. Carbon, Nitrogen and oxygen atoms are colored grey, blue and red respectively. Surface of actin is drawn to show the occluded and exposed state of a cryptic site in the unbound and ligand-bound structures of actin and is superposed with its unordered carboxy-terminus, protrusion of which into the site makes the site cryptic. (A) Occluded cryptic site due to the protrusion of unordered carboxy-terminus (shown as yellow cartoon) in the unbound structure of actin (yellow, (1RDW(X)) is shown. (B) Binding of ligand tetramethylrhodamine-5-maleimide (TMR) (Ligand ID RHO, grey) at the exposed cryptic site with missing carboxy-terminus is shown for actin (smudge, 1J6Z(A)). (C) Binding of ligand ethylene glycol (Ligand ID EDO, grey/darkgrey) at the exposed cryptic site with ordered carboxy-terminus (shown as blue/darkblue cartoon) is shown for actin (slate, 4B1V(A),4B1V(B)). (D) Schematic representation of the interactions made by ligands tetramethylrhodamine-5-maleimide (TMR) (Ligand ID RHO) and ethylene glycol (Ligand ID EDO), with actin. Actin residue identifiers (residue name, number, chain ID) and ligand identifiers (Ligand ID, number, chain ID) along with atom names of bound ligands are indicated. Hydrogen bond length is given in Å. The red circles and ellipses indicate actin residues that are in equivalent 3D positions when complexes of actin with ligands RHO (1J6Z(A)) and EDO (4B1V(A),4B1V(B)) are superposed thereby highlighting the actin-RHO and actin-EDO interactions that are common at the exposed cryptic site of actin.

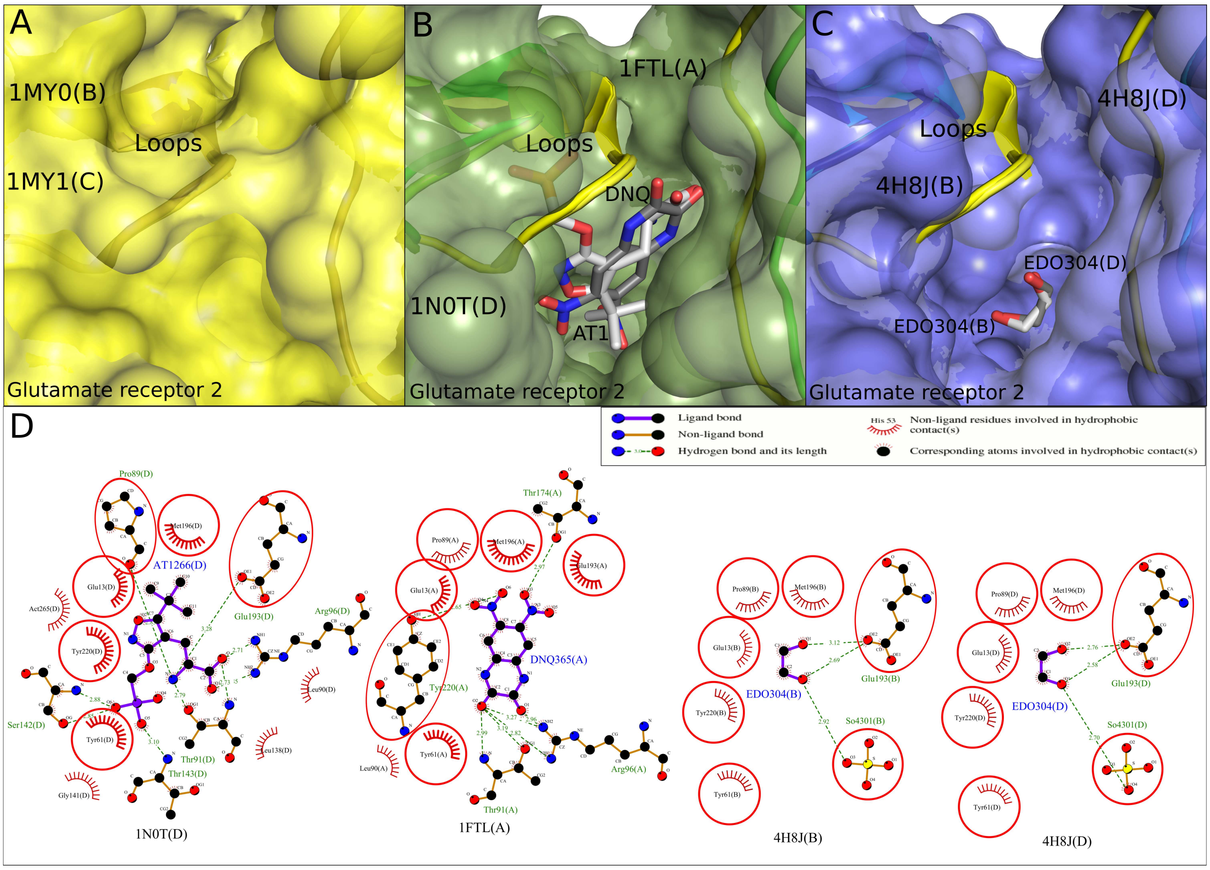


Figure S6: **Ethylene glycol identifies a cryptic site in Glutamate receptor 2.** Bound ligands are shown in a stick model. Carbon, Nitrogen and oxygen atoms are colored grey/darkgrey, blue and red respectively. Surface of glutamate receptor 2 is drawn to show the occluded and exposed state of a cryptic site in the unbound and ligand-bound structures of glutamate receptor 2 and is superposed with its loops, protrusion of which into the site makes the site cryptic. (A) Occluded cryptic site due to protrusion of loops (shown as yellow cartoon) in the unbound structure of glutamate receptor 2 (yellow, (1RDW(X)) is shown. (B) Binding of ligands ATPO (Ligand ID AT1, grey), DNQX (Ligand ID DNQ, darkgrey) at the exposed cryptic site with displaced loops (shown as green/darkgreen cartoon) is shown for glutamate receptor 2 (smudge, 1N0T(D),1FTL(A)). (C) Binding of ligand ethylene glycol (Ligand ID EDO, grey/darkgrey) at the exposed cryptic site with displaced loops (shown as blue/darkblue cartoon) is shown for glutamate receptor 2 (slate, 4H8J(B),4H8J(D)). (D) Schematic representation of the interactions made by ligands ATPO (Ligand ID AT1), DNQX (Ligand ID DNQ) and ethylene glycol (Ligand ID EDO), with glutamate receptor 2. Glutamate receptor 2 residue identifiers (residue name, number, chain ID) and ligand identifiers (Ligand ID, number, chain ID) along with atom names of bound ligands are indicated. Hydrogen bond length is given in Å. The red circles and ellipses indicate Glutamate receptor 2 residues that are in equivalent 3D positions when complexes of Glutamate receptor 2 with ligands AT1 (1N0T(D)), DNQ (1FTL(A)) and EDO (4H8J(B),4H8J(D)) are superposed thereby highlighting the interactions of Glutamate receptor 2 with ligands AT1,DNQ and EDO that are common at the exposed cryptic site of Glutamate receptor 2.
